# Mannosidases IA, IB and IC are in segregated vesicular structures and involved in both glycoprotein quality control and maturation

**DOI:** 10.64898/2026.06.07.730669

**Authors:** Haddas Saad, Marina Shenkman, Edward Avezov, Isam Khalaila, Gerardo Z. Lederkremer

## Abstract

N-linked glycoprotein processing critically depends on the trimming of α-1,2 mannose residues, a key step required both for glycoprotein maturation along the secretory pathway and for targeting defective glycoproteins to endoplasmic reticulum-associated degradation (ERAD). Mammalian cells express seven Class I α-1,2 mannosidases, yet their individual roles remain poorly defined, particularly for ManIA, ManIB, and ManIC, which were originally considered Golgi-resident maturation enzymes. Here, we re-evaluated the subcellular localization and functional contributions of these three mannosidases to glycoprotein quality control and maturation. We found that ManIA, ManIB, and ManIC localize predominantly to quality control vesicles (QCVs), previously identified by our group, whereas only ManIC displays a substantial Golgi population. Surprisingly, each enzyme is confined to a different vesicular population. All three enzymes promote ERAD targeting of misfolded model glycoproteins, albeit with different substrate preferences. In addition, they redundantly support the maturation and cell-surface delivery of a model glycoprotein. Most strikingly, in vitro analyses revealed that ManIA, ManIB, and ManIC preferentially trim a properly folded model glycoprotein rather than its denatured form. This is the opposite of the substrate preference that we previously observed for ERManI, EDEM1, and EDEM2. These findings support a model in which ERManI and the EDEMs selectively process misfolded glycoproteins to promote their recognition by the proximal lectin OS-9 and subsequent ERAD. In contrast, ManIA, ManIB and ManIC can slowly process misfolded glycoproteins but act rapidly on properly folded glycoprotein molecules, releasing them from ER-Golgi lectin-mediated retention or retrieval pathways, thereby promoting forward trafficking and maturation in the Golgi.

## Introduction

The most common posttranslational modification on secretory and membrane proteins in mammalian cells is N-glycosylation, the addition of a glycan precursor containing 14 sugar units to an asparagine residue during protein translocation into the endoplasmic reticulum (ER). Most of these N-glycans then undergo processing, which involves removal of glucose and mannose residues and addition in the Golgi apparatus of other sugars such as GlcNAc, Gal and sialic acid to render complex-type sugar chains (1, 2). Removal of α-1,2 linked mannose residues is required for this addition and obtention of complex-type chains. Interestingly, if this removal takes place too early, before exit of the glycoprotein from the ER, it leads to its targeting to ER-associated degradation (ERAD) (3). The longer it takes for a glycoprotein to exit the ER (e.g. for misfolded molecules), the higher the probability of undergoing this trimming and being sent to ERAD (4, 5). The α-1,2 mannosidases that have been implicated in the pre-Golgi removal are ER mannosidase I (ERManI/MAN1B1) (6) and three ER-degradation enhancing mannosidase-like proteins (EDEM1-3) (7). In line with this, it was shown that ERManI, EDEM1 and EDEM2 have preference for unfolded/misfolded glycoprotein molecules rather than properly folded ones (8, 9). The removal of the α-1,2 Man residues leads to recognition of the glycoprotein molecules by the ERAD-complex-associated lectins OS-9 and XTP3-B (4, 10, 11). Before their removal, the folding-sensor UGGT1 recognizes unfolded states of the glycoprotein molecules and adds one Glc residue to the glycans (12), allowing recognition by the lectin/chaperones calnexin (CNX) or calreticulin (CRT) for another folding attempt (1, 13). Thus, the extensive mannose trimming serves three aims: First, the removal of a terminal mannose that is the Glc acceptor of UGGT1, precluding re-entry of the glycoprotein into the CNX folding cycle (14). Second, the lectin OS-9 or its functional homolog XTP-3B, which target glycoproteins to ERAD, bind with high affinity to the trimmed M5-6 structures and they cannot bind the untrimmed glycan (4, 15). Third, the trimmed species do not bind the lectins VIPL, VIP36 and ERGIC53, and are therefore selected against in ER-Golgi transport (16, 17). For molecules that fold correctly and exit to the Golgi, the removal of α-1,2 Man residues for glycoprotein “maturation” is done by *Class I* “Golgi” α-1,2 mannosidases, ManIA (MAN1A1), IB (MAN1A2) and IC (MAN1C1) (18). It is unknown why 3 different mannosidases with similar activities (19, 20) are needed. Overexpression of any of these three mannosidases was shown to accelerate ERAD of misfolded human α1-antitrypsin variant null Hong Kong (NHK) (21), but overexpression of the mannosidases might increase their ER levels during their biosynthesis causing premature N-glycan trimming in the ER. However, ManIA was shown to indeed be involved in targeting misfolded glycoprotein molecules to ERAD (22, 23). Furthermore, ManIA is not located in the Golgi but instead in specialized quality control vesicles (QCVs) (23). We recently observed that ManIB and ManIC also appear partially in a vesicular pattern (24). Here we find that ManIB and IC, besides participating in glycoprotein maturation, are also involved in glycoprotein quality control and are present in QCVs. Surprisingly, each enzyme resides in a different vesicular population.

## Materials and Methods

### Materials

Lipofectamine 2000 transfection reagent was from Invitrogen (Cat#52758). cOmplete Protease Inhibitor Cocktail was from Roche (Cat#11697498001). Promix cell labeling mix ([^35^S]Met plus [^35^S] Cys, 1000 Ci/mmol) was from PerkinElmer Life Sciences. Streptavidin-Agarose was from Millipore (S1638), Sulfo succinimidyl-6-biotin (Cat#935409), OptiPrep Density Gradient Medium (Cat#D1556), BrefeldinA (Cat#B6542) and other common reagents were from Sigma-Aldrich.

### Plasmids and constructs

ManIA-HA, ManIB-HA and ManIC-HA in the pMH expression vector (Roche Diagnostics, Basel, Switzerland) were described before (24). ManIA-mCherry was described before (23). ManIB cDNA was subcloned into eGFP-N1 expression vector (Clonetech) with a flexible linker (SGGGGS), using Sal1 and HindIII. ManIC-GFP was constructed similarly to ManIB construct but using ManIC cDNA. ERGIC53-GFP and GalT-YFP (first 60 amino acids of β1,3-galactosyltransferase linked to YFP) were used before (4). GFP-Rab1a and Sar1[T39N]-GFP were previously used(4). Sar1[T39N] is in an IRES-GFP vector, which allows observation of Sar1[T39N] expression in live cell imaging without direct fusion of a fluorescent protein to it. BACE476-HA and NHK-HA were described before (25). H2a subcloned in pCDNA1 and the pSUPER vector carrying an shRNA for ManIA were described before (23). H1 subcloned in pCDNA1 was described before (26). Predesigned shRNAs for ManIB (TRCN000049629, TRCN000049630, TRCN000049631, TRCN000049632),ManIC (TRCN000049639, TRCN000049640, TRCN000049641, TRCN000049642) EDEM2 (TRCN000369425, TRCN000342980, TRCN000298248, TRCN000055969, TRCN000286597) and EDEM3 (TRCN0000156214, TRCN0000151874, TRCN0000151952) were from Sigma-Aldrich. The pSUPER vector carrying an shRNA for human ERManI, pSUPER encoding anti–human EDEM1 shRNA, pSUPER encoding anti-lacZ shRNA, and S-tagged OS-9.1 and OS-9.2 were those described before (8). LentiCRISPR v2 was from addgene (Plasmid #52961). Guide RNAs (gRNAs) targeting human ManIA (CACCGGGATCCGCGAAAACCACGAG), ManIB (CACCGCCACATGTAGATGCCGGTAA) and ManIC (CACCGAGCTATAAGCGTTATGCAAT) were from Sigma-Aldrich. VSV-G and DR8-2 plasmids were a kind gift from Eran Bacharach (TAU).

### Antibodies

Mouse anti-HA was from Biolegend (Cat#901514), rabbit anti-calnexin (Cat#C4731), rabbit anti-EEA1 (Cat#E3906), mouse anti-GAPDH (Cat#G8795) and mouse anti-actin (Cat#A3853) were from Sigma-Aldrich. Rabbit anti-Cab45 was the one used previously (27). Rabbit anti-ManIA (Cat#M3694), anti-ManIB (Cat#SAB2101422) and anti-ManIC (Cat#SAB4200243) were from Sigma-Aldrich. Mouse anti-ERManI was previously used (4). Rabbit anti-EDEM1 (Cat#E8406), anti-EDEM2 (Cat#E1782) and anti-EDEM3 (Cat#E8781) were from Sigma-Aldrich. Rabbit polyclonal anti-H2a antibody was the one used in previous studies (8). Rabbit polyclonal anti-GM130 was from BioLegend, anti-S-tag was from Novagen. Antibodies specific for peptides corresponding to the carboxyl termini of H1 were the ones used in a previous study (26). Goat anti-mouse IgG-HRP (Cat#111-035-144) and goat anti-rabbit IgG-HRP (Cat#115-035-166) were from Jackson-Immuno Research Labs.

### Cell culture, media and transfections

NIH 3T3 and HEK 293 cells were grown in DMEM supplemented with 10% bovine calf serum at 37°C under 5% CO_2_. Transfections of NIH 3T3 cells were carried out using Lipofectamine 2000 transfection reagent. Transfection of HEK 293 cells was performed using the calcium phosphate method.

### Production of ManIA, ManIB and ManIC CRISPR/Cas9 knockout cell lines

Three constructs with different guide RNAs (gRNAs) targeting each of the human ManIA, ManIB and ManIC genes, in a vector containing the Cas9 gene and a selection marker, were used to generate the knockout cell lines. The three constructs were co-transfected with VSV-G and DR8-2 packaging plasmids into HEK 293T cells. 48 hours post transfection, the viral-like particles were collected and used to infect HEK 293 cells. Following 12 days of Puromycin selection (0.03µg/ml), the cells were serially diluted. Cells from selected clones were lysed and the knockout of ManIA, ManIB or ManIC was confirmed by immunoblotting.

### Immunofluorescence Microscopy

Immunofluorescence was performed as described previously (23). Briefly, for live cell microscopy, cells were transfected and grown for 24h on glass bottom 35mm plates. Images and time-lapse videos were captured at 37°C under 5% CO_2_ using a Leica SP8 laser scanning confocal microscope. For fixation purposes, cells grown for 24h after transfection on coverslips in 24-well plates were fixed with 3% paraformaldehyde for 30 min, incubated with 50mM glycine in PBS, and left unpermeabilized or permeabilized with 0.5% Triton X-100. After blocking with normal goat IgG in PBS/2% BSA, they were incubated with a primary antibody for 60 min, washed and incubated for 30 min with a secondary antibody, followed by washes. Specimens were observed using a Leica SP8 laser scanning confocal microscope or LSM 510 meta microscope. ImageJ was used to quantify fluorescence intensity and to calculate Manders’ coefficients for colocalization studies.

### Iodixanol equilibrium sedimentation gradient

Iodixanol gradients were performed similarly to previously described (27). Briefly, HEK 293 cells transfected with plasmids as mentioned in the results were washed with PBS and resuspended in a homogenization buffer (0.25M Sucrose, and 10mM HEPES pH7.4). The cells were passed through a 21G needle 5 times before homogenization in a Dounce homogenizer (low clearance pestle, 30 strokes). The homogenates were centrifuged at 1000xg for 10 minutes at 4 °C to remove nuclei and cell debris and the supernatants were loaded on top of an iodixanol gradient (10 to 34%). The gradients were ultra-centrifuged at 24,000 rpm (98,500g, Beckman SW41 rotor) at 4 °C for 16 hours. 1ml gradient fractions were collected from top to bottom and run on SDS-PAGE.

### Velocity sedimentation gradient

Velocity sedimentation gradients were performed as previously described (23). In brief, post-nuclear supernatants were loaded on glycerol gradients [9.5 ml 10–30% glycerol on top of 0.5ml 50% sucrose, in10mM Hepes (pH 7.4), 0.15 M NaCl, 1 mM EGTA, and 0.1 mM MgCl2]. The gradients were ultracentrifuged at 218,309×g in a Beckman SW41 rotor, for 45 min at 4 °C. Ten fractions were collected from top to bottom and subjected to immunoblotting.

### Transmission Electron Microscopy (TEM)

Aliquots from the first eight fractions collected from the velocity gradient were applied to freshly glow-discharged 200-mesh Formvar-coated copper grids and allowed to adsorb for 1 minute. Excess liquid was removed via filter paper wicking, and the grids were negatively stained with 1% uranyl acetate for 30 seconds before being air-dried at room temperature. The samples were visualized using a JEM-1400 Plus transmission electron microscope (JEOL, Tokyo, Japan) operated at an accelerating voltage of 120kV. Images were captured using a Gatan OneView digital camera at magnifications ranging from 40Kx to 250Kx. The procedure and imaging was done with the help of Vered Holdengreber.

### Dynamic Light Scattering (DLS)

The hydrodynamic diameter and size distribution of the particles in the first eight velocity gradient fractions were determined by Dynamic Light Scattering (DLS). Measurements were performed using a Malvern Zetasizer Nano ZS equipped with a 633 nm He-Ne laser. Each fraction was measured at 25°C using a disposable micro-cuvette. Samples were equilibrated for 120 seconds prior to data collection. For each fraction, three independent measurements were recorded, with each measurement consisting of 10 to 15 individual runs. The scattering intensity was captured at an angle of 173° (backscatter detection).

The translation diffusion coefficient was used to calculate the mean hydrodynamic diameter (Z-average) and the polydispersity index (PdI) via the Stokes-Einstein equation:

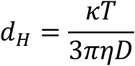

(Where d_H_ is the hydrodynamic diameter, κ is the Boltzmann constant, T is absolute temperature, η is solvent viscosity, and D is the diffusion coefficient). Data were processed using Zetasizer Software v7.13, results are expressed as the mean ±SD.

The measurements were performed at the “Tel Aviv University Center for Nanoscience and Nanotechnology” with the advice of Dan Peer and the assistance of Nicole Gorokhovsky.

### Pulse-chase analysis

Pulse-chase analysis was performed similarly to previously described (23). In brief, HEK293 cells were incubated for 30 min in Cys-free medium and divided equally into Eppendorf tubes. Cells were labeled with [^35^S]Cys for 20 min, rinsed and transferred to full medium for indicated chase periods followed by lysis and immunoprecipitation with anti-H2a or anti-HA antibodies. SDS-PAGE was performed on 12% gels imaged by fluorography and analyzed by phosphorimager quantitation.

### Immunoprecipitation and immunoblotting

Cell lysis and immunoprecipitation were done as described before (11). Briefly, cells were lysed (1% Triton X-100 and 0.5% sodium deoxycholate in PBS) in the presence of cOmplete Protease Inhibitor Cocktail for 30 min on ice. After pelleting of nuclei, lysates were immunoprecipitated with protein A-Sepharose and the appropriate antibody, incubated at 4°C with rotation for 16 h, followed by three washes with 0.5% Triton X-100, 0.25% sodium deoxycholate and 0.5% SDS in PBS and once with PBS. Samples were boiled in sample buffer for 5 minutes and run on SDS-PAGE under reducing conditions. Transfer to a nitrocellulose membrane, immunoblotting, detection by ECL, and quantitation in a Bio-Rad ChemiDoc XRS Imaging System (Hercules, CA) were done as described previously. ImageJ software was used for the quantitation.

### Cell surface biotinylation

HEK 293 cells were incubated with Sulfo-succinimidyl-6-biotin hexanoate (0.02 mg/mL in PBS) for 45 minutes at 4°C with constant rocking. The biotinylation reaction was quenched by incubation with 50mM NH_4_Cl in PBS for 10 minutes at 4°C. The cell lysates were subjected to precipitation using streptavidin beads with overnight incubation at 4°C. The beads were washed, and the captured biotinylated fraction was eluted for analysis via SDS-PAGE.

### N-glycan analysis

N-glycan analysis was performed as previously described (8). In brief, to isolate HA-tagged ManIA, ManIB and ManIC, they were expressed in subconfluent (90%) monolayers of HEK 293 cells. 24h post transfection, the cells were lysed in 1% NP-40, CaCl_2_ 5mM, PIPES 0.1M pH7.5, and protease inhibitor cocktail for 30 min on ice, and debris and nuclei were pelleted in a microfuge for 30 min at 4°C. The samples were immunoprecipitated with anti-HA antibody and anti-mouse IgG agarose beads overnight at 4°C.

The beads containing immunoprecipitated mannosidases were then washed twice with reaction buffer (0.2% NP-40, 5 mM CaCl_2_, PIPES 0.1M pH7.5), followed by overnight incubation with M. rosenbergii high-density lipoprotein (HDL) containing vitellogenin in its native or denatured form in the reaction buffer at 37 °C. HDL was purified from a mix of hemolymph, withdrawn from M. rosenbergii secondary vitellogenic females, and 7% ethylenediaminetetraacetic acid in 1:1 ratio with the addition of 2mM phenylmethylsulphonyl fluoride (Sigma). The haemolymph solution was then mixed with a saturated solution of sodium bromide to a density of 1.22gml−1. HDL containing the egg yolk protein vitellogenin was isolated by ultracentrifugation at 100,000 g for 48 h. The upper golden fraction (HDL) was collected and dialyzed against 20mM Tris buffer pH 8.0. To denature vitellogenin, it was treated with 6M guanidine chloride and 20mM dithiothreitol (DTT) for 6h at room temperature (RT), followed by addition of 200mM iodoacetamide for 30min at RT and overnight dialysis against reaction buffer at 4°C.

After overnight treatment with the immunoprecipitated mannosidases, vitellogenin was reduced (55mM DTT, 70°C), alkylated (100mM iodoacetamide), and resolved on 7% SDS-PAGE. The 89kDa bands were excised, destained, and dehydrated via SpeedVac. N-glycans were released by Endo-H digestion and extracted using water and ACN. Finally, glycans were labelled with 2-aminobenzamide (2-AB), analyzed by normal-phase High Performance Liquid Chromatography (HPLC), and quantified using ImageJ. The separation of N-linked oligosaccharides by HPLC was performed by Dr. Isam Khalaila (BGU, Israel) or by Yehudit Amor, Savyon Diagnostics Ltd, Israel.

### Statistical analysis

The results are expressed as average ±SD or mean ±SEM as indicated. Student’s t-test (two-tailed) was used to compare the averages of two groups. Statistical significance was determined at P<0.05 (*), P<0.01 (**), P<0.001 (***), P<0.0001 (****).

## Results

### Class I α-1,2 mannosidases IB and IC reside in segregated vesicles with only a fraction of their populations in the Golgi

Earlier studies provide conflicting observations concerning the localization of Golgi class I α-1,2 mannosidases. The unexpected finding that ERManI and ManIA are in QCVs rather than in the ER or the Golgi (4, 23), and our recent observation that ManIB and ManIC are also partially located in vesicular structures (24), prompted us to further investigate the subcellular location and roles of these enzymes. We examined the localization of the endogenous and exogenous ManIB and ManIC in NIH 3T3 cells, using immunofluorescence. The cells were transfected with either β-1,3 galactosyltransferase linked to YFP (GalT-YFP) as a Golgi marker, or with ERGIC53-GFP as an ERGIC marker, with or without co-transfection with ManIB-HA or ManIC-HA. 24 hours post transfection the cells were fixed and subjected to immunofluorescent staining. Surprisingly, in most cells, endogenous ManIB appeared in a punctate and patchy pattern, colocalizing only to a minor extent with GalT-YFP (Fig. 1A, K). Similarly, only a minor population of ManIB colocalized with ERGIC53-GFP (Fig. 1B, L). ManIC also appeared in a punctate and patchy pattern, but with a larger population in the juxtanuclear region colocalizing with GalT-YFP and ERGIC53-GFP (Fig. 1C, D, K, L). Exogenous ManIB-HA and ManIC-HA showed a similar pattern to that seen with endogenous ManIB and ManIC (Fig. 1E-H, K, L). In both cases, the enzymes appeared in a punctate pattern, only partially located in the juxtanuclear region with colocalization with the ERGIC and Golgi markers. This juxtanuclear population was more conspicuous for ManIC (Fig. 1, arrows). To analyze whether the punctate structures might derive from the Golgi, cells transfected with GalT-YFP and either ManIB-HA or ManIC-HA, were treated with Brefeldin A (BFA) (Fig 1I, J). BFA causes Golgi complex collapse into the ER (28), as can be seen by partial dispersion of GalT-YFP. In contrast, ManIB and ManIC remained mostly in a similar punctate pattern (through BFA insensitivity or possible rearrangements), and there was some dispersion for ManIC (Fig. 1I, J). The extent of colocalization with GalT-YFP was not significantly changed by BFA treatment, remaining low for ManIB and higher for ManIC (Fig. 1K). To observe the subcellular localization of ManIB and ManIC in live cells and dispel concerns that cell fixation and permeabilization could have affected ManIB and ManIC localization (29), we used fusion proteins of these mannosidases with the fluorescent protein GFP. This also enabled us to compare their localization to that of ManIA linked to mCherry in live cells. Live cell images of NIH 3T3 cells co-expressing ManIA-mCherry and ManIB or ManIC fused to GFP, show part of the population of ManIB and ManIC in a punctate pattern with partial colocalization with ManIA (Fig. 1M-O). In addition, there were juxtanuclear accumulations of ManIB-GFP and especially of ManIC-GFP, which did not colocalize with ManIA-cherry. Overall, the colocalization with ManIA-cherry was much lower for ManIC-GFP than for ManIB-GFP (Fig. 1O).

**Figure 1.**
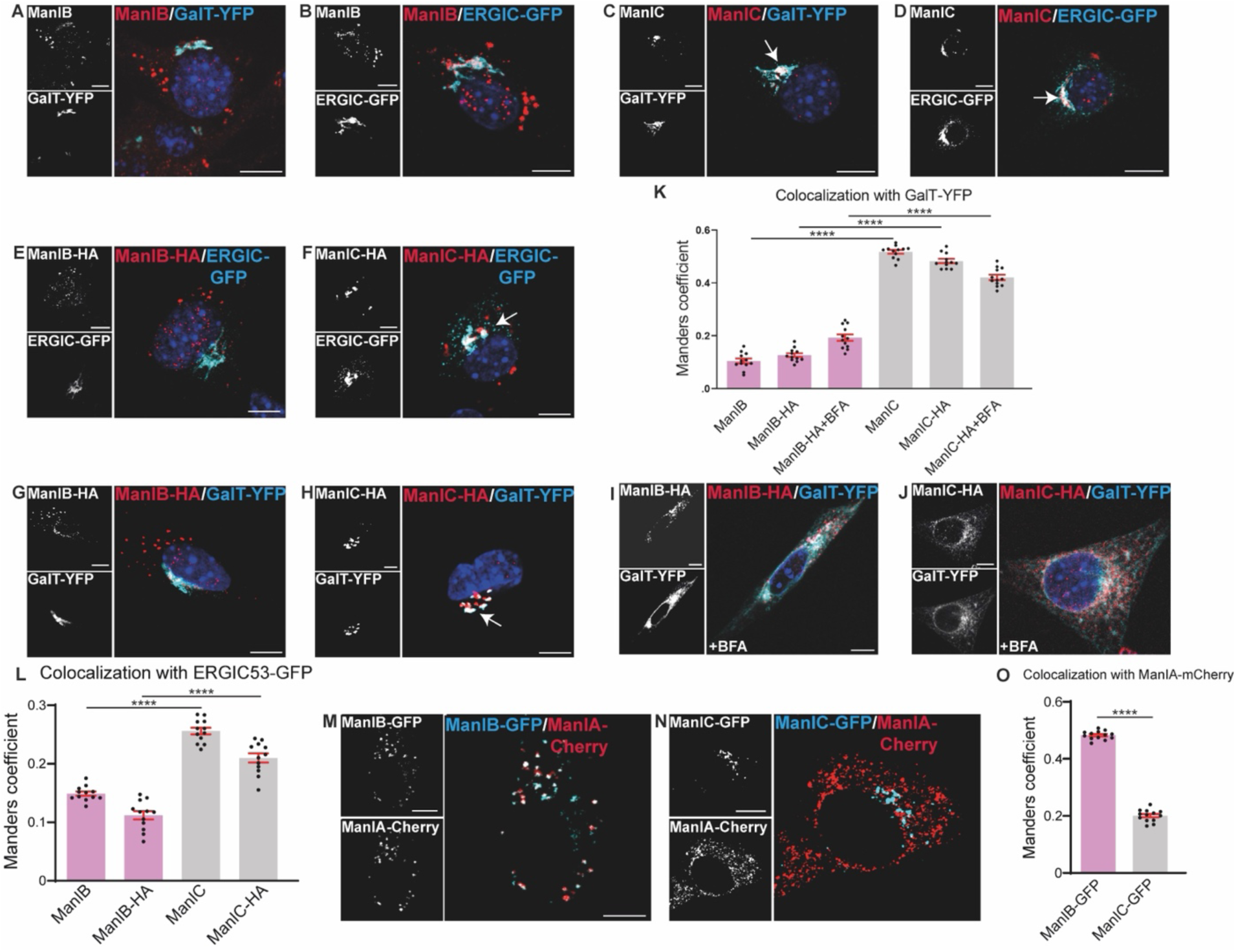
Endogenous and exogenous class I α1,2 mannosidases IB and IC appear in a vesicular pattern, only partially colocalizing with the Golgi and the ERGIC. **(A-L)** NIH 3T3 cells were transfected with the Golgi marker GalT-YFP or the ERGIC marker ERGIC53-GFP and ManIB-HA or ManIC-HA, as indicated. Cells were fixed and subjected to immunofluorescent staining with rabbit anti-ManIB **(A, B)** or anti-ManIC **(C, D)** and Dylight 650-conjugated goat anti-rabbit IgG to detect the endogenous mannosidases, or with mouse anti-HA and Dylight 649-conjugated goat anti-mouse IgG **(E-J).** At 24 h post-transfection, some samples were treated with BFA (5 μg/ml for 30 min) **(I, J)**. For better visualization, merged confocal images were pseudo-colored red (ManIB and ManIC) and cyan (GalT-YFP and ERGIC53-GFP). The arrows indicate the juxtanuclear accumulation of ManIC. **(K-L)** The graphs represent the colocalization of endogenous and HA-tagged ManIB and ManIC with GalT-YFP **(K)** or ERGIC-GFP **(L).** Representative results of three independent experiments are presented (n=12 for each protein). **(M-O)** Live cell imaging of NIH 3T3 cells co-expressing ManIA-mCherry and ManIB-GFP **(M)** or ManIC-GFP **(N)**. For better visualization, merged confocal images were pseudo-colored cyan (ManIB-GFP and ManIC-GFP) and red (ManIA-mCherry). Representative results of three independent experiments are presented. **(O)** The graph represents the colocalization of ManIB-GFP or ManIC-GFP with ManIA-Cherry (n=13 for each protein).

### ManIA, ManIB and ManIC reside in different types of vesicles

The partial colocalization of ManIB and ManIC puncta with ManIA punctate structures, their lack of colocalization with ERGIC or Golgi markers and their insensitivity to BFA suggested that they may localize to vesicular structures that we termed quality control vesicles (QCVs) (23). We had observed that QCVs partially colocalize with the small GTPase Rab1a (4), thus we examined the localization of ManIB and ManIC relative to GFP-Rab1a (Fig. 2A-C). Rab1a has been associated with COPII carrier formation and targeting. In addition to its presence in QCVs, Rab1a accompanies secretory vesicles toward the Golgi and participates in their tethering and subsequent delivery to the Golgi membrane(30). Therefore, it can be observed in both vesicular and Golgi structures (Fig. 2A, B). Co-expression of ManIB-mCherry with GFP-Rab1a revealed colocalization of the two proteins in a population of cytoplasmic vesicles (Fig. 2A, C). In contrast, ManIC-mCherry showed colocalization with GFP-Rab1a predominantly in the Golgi region (Fig. 2B, C).

**Figure 2.**
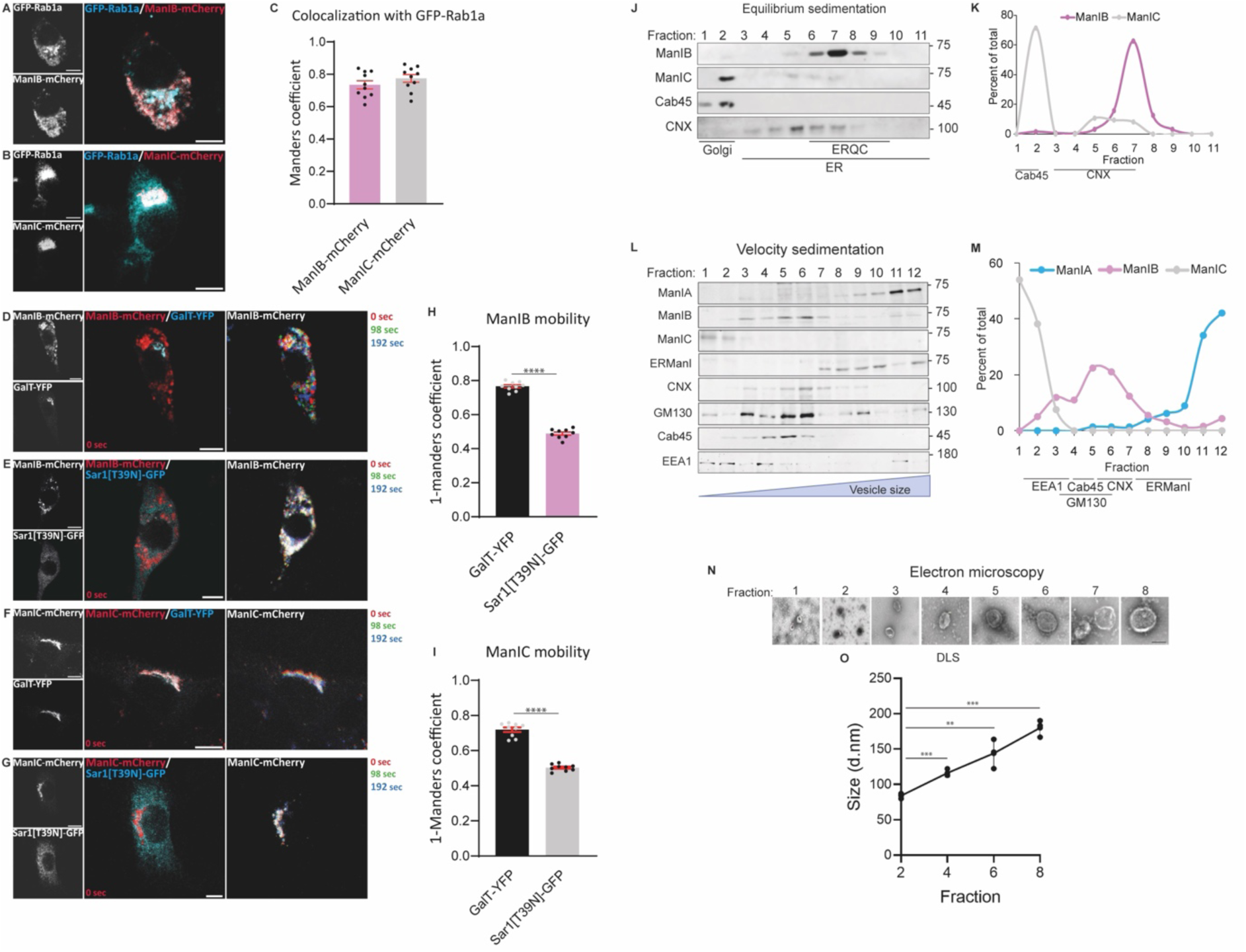
ManIA, ManIB and ManIC reside in mobile quality control vesicles of different sizes. **(A-C)** NIH 3T3 cells coexpressing GFP-Rab1a and ManIB-mCherry (a) or ManIC-mCherry (B). At 24h post-transfection, the cells were fixed with 3% PFA. For better visualization, merged confocal images were pseudo-colored red (ManIB and ManIC) and cyan (GFP-Rab1a). Representative results of three independent experiments are presented (n=10 for each protein), Bars=10µm. The graph represents the colocalization of ManIB and ManIC with GFP-Rab1a, as inferred by Manders’ coefficient**. (D-I)** Live-cell imaging of NIH 3T3 cells expressing ManIB-mCherry (D, E) or ManIC-mCherry (F, G) together with GalT-YFP (D, F) or dominant-negative Sar1[T39N]-GFP (E, G). In the right panel only the mCherry channel is shown and three time points were selected, pseudocolored as indicated, and superimposed to visualize vesicle movement. Representative results of three independent experiments are presented (n=9 for each), Bars=10µm. ManIB-mCherry (H) and ManIC-mCherry (I) mobility was quantified using the Manders’ coefficient by comparing localization at 98 sec with that at 0 sec. Mobility is presented as 1 – Manders’ coefficient, with lower values representing reduced mobility. **(J)** HEK 293 cells were homogenized, and the homogenate was loaded on top of an iodixanol gradient (10%-34%). The gradients were ultracentrifuged at 24,000 rpm (SW41 rotor) at 4C for 16 hours. Eleven fractions were collected from the top to the bottom (1-11 is from the lightest to the heaviest fraction), run on 12% SDS-PAGE and immunoblotted with anti-ManIB or anti-ManIC, anti-CNX anti-Cab45 (Golgi marker) antibodies. **(K)** The intensity of each band was acquired and quantified by ImageJ. The results shown are representative of three independent experiments. **(L)** Post-nuclear supernatants of HEK 293 cells were loaded on 10%–30% glycerol gradients on top of a 50% sucrose cushion, fractionated to 10 fractions top to bottom, and immunoblotted with rabbit anti-ManIA, anti-ManIB and anti-ManIC, mouse anti-ERManI, rabbit anti-CNX, anti-GM130, anti-Cab45 and anti-EEA1. **(M)** The graph shows quantitation of the immunoblot. The results shown are representative of three independent experiments. **(N)** Samples of fractions 1-8, taken from the velocity sedimentation gradient presented in (l), were viewed using a JEM-1400plus transmission electron microscope. Bar= 120 nM **(O)** The diameter of vesicles in each fraction was quantified using dynamic light scattering (DLS), resulting in a median diameter of 84.07±3,75 in fraction 2 up to 180.03±12.04 in fraction 8. Median of three independent experiments ±SEM.

We had also observed previously that QCVs are highly mobile and that this mobility is dependent on COPII. To examine the mobility of ManIB- and ManIC-containing vesicles, we performed live-cell imaging in the absence or presence of dominant-negative Sar1 (Sar1[T39N]-GFP), which interferes with COPII-dependent ER export (31) (Fig. 2D-I and supplemental videos S1-S4). The mobility of ManIB-mCherry and ManIC-mCherry was assessed by comparing their localization at the start of the live imaging (0 sec) with that after two different time frames (Fig. 2D-G, right panels). These time frames were overlaid and pseudocolored and all three colors can be observed where the vesicles are mobile, but much less mobile structures containing ManIC-mCherry in the Golgi region appear white. The mobility is considerably reduced in the presence of dominant-negative Sar1. The change in mobility was quantified by the reduction in colocalization after 98 sec, using the Manders’ coefficient and expressing mobility as 1 – Manders’ coefficient. Because Manders’ coefficient quantifies spatial overlap rather than displacement, its complement was used to express the loss of overlap with the initial distribution as a mobility index, such that increasing values correspond to increasing spatial redistribution over time. In the presence of dominant-negative Sar1, the mobility of both ManIB and ManIC was significantly reduced compared with the high mobility in cells co-expressing GalT-YFP as a control (Fig. 2H, I). These findings indicate that ManIB- and ManIC-containing vesicles are highly dynamic and sensitive to disruption of Sar1-dependent trafficking, pointing to COPII machinery involvement, as had been observed for QCVs(4).

We also analyzed the localization of endogenous ManIB and ManIC using a different approach, by performing iodixanol equilibrium sedimentation gradients optimized for separation of the Golgi complex and ER subdomains based on their density. ManIB migrated in equilibrium sedimentation at a peak at slightly denser fractions compared to the ER marker CNX (Fig. 2J, K). This was consistent with its presence in the fluorescence images in mainly a punctate pattern with only a minor population colocalizing with a Golgi marker (Fig. 1) and similar to what we had observed for ERManI and ManIA in QCVs (23). In contrast, ManIC migrated in equilibrium sedimentation mostly at a Golgi-like density, comigrating with the Golgi protein Cab45, with only a smaller fraction at ER-like densities (Fig. 2J, K), in line with its significant colocalization with the Golgi marker, and a smaller population in a punctate pattern in the fluorescence images (Fig. 1).

Next, we performed velocity sedimentation experiments, which provide a separation according to the sedimentation rate instead of density. This technique, which is a modification of a protocol used for the separation of synaptic vesicles on glycerol gradients, allowed an improved resolving capability for QCVs containing ManIA (23). These gradients separate vesicles and organelle-derived microsomes of different sizes without differentiating their origin. We compared the migration of ManIA, ManIB, ManIC and also ERManI. Endogenous ManIA migrated mostly in the heaviest fractions 11-12 (Fig. 2L, M). Endogenous ManIB migrated mostly in the middle fractions, similar to CNX, Cab45 and most of GM130 (Fig. 2L, M). Only a minor population of ManIB appeared in heavier fractions, in the same region where ManIA migrated. ManIC, on the other hand, migrated only in the light fractions, suggesting its presence in very small vesicles, similar to the ones derived from the endosomes (EEA1) (Fig. 2L, M). It is important to note that in this assay, ManIA and ManIC vesicles appear to be largely devoid of Golgi or ER markers, suggesting that these vesicles do not originate from either organelle (Fig. 2L). ManIB-containing vesicles, in contrast, comigrate with both ER and Golgi markers. Since velocity sedimentation separates structures based primarily on their physical properties and does not establish their subcellular origin, this comigration does not necessarily indicate that ManIB-containing vesicles derive from either organelle. Surprisingly, despite the high level of homology between the three mannosidases, they appear almost entirely separated in different fractions in the velocity gradient, indicating that they are segregated in different types of vesicles.

The velocity sedimentation gradients separate vesicles of different sizes, the smallest vesicles appearing in the top fraction 1 and the size of the vesicles increases when moving down the gradient. However, vesicle shape can significantly affect the distribution. We examined the morphological characteristics of the vesicles in the first eight fractions collected from the velocity gradient using transmission electron microscopy (TEM), which confirmed that the vesicles are indeed distributed along the gradients according to their sizes (Fig. 2N). However, there appears to be some heterogeneity, with a range of sizes in each fraction. To analyze quantitatively the distribution of these vesicles, we performed dynamic light scattering (DLS) analysis, which can provide statistically significant information regarding the distribution of the sizes of the vesicles. The results show that indeed the vesicles increase linearly in size as we move along the gradient, with a median diameter of 84.07±3,75 nm in fraction 2 up to 180.03±12.04 nm in fraction 8 (Fig. 2O).

### ManIA participates in targeting to ERAD of H2a, whereas ManIB and ManIC do not

Although the Golgi α-1,2 mannosidases IA, IB and IC play a central role in glycoprotein maturation, the level of redundancy or specificity in their roles remained unclear. It has previously been reported that the overexpression of any of the three Class I α-1,2 mannosidases accelerated ERAD of misfolded human α1-antitrypsin variant null Hong Kong (NHK) (21). One plausible explanation is that the trimming of precursor N-glycans in the ER is increased unnaturally due to the overexpression of these mannosidases, which could increase their ER levels during their biosynthesis and trafficking. Alternatively, as NHK was shown to partially exit the ER, being then retrieved, NHK molecules could travel to the Golgi where they could be targeted by those enzymes. In a previous study we tested the effect of overexpression of ManIA on the fate of another model ERAD substrate, unassembled CD3δ chain of the T cell receptor, which was much increased by ManIA overexpression (23). The same study showed that the overexpression of ManIA also increased the degradation of another established ERAD substrate, the uncleaved precursor of the asialoglycoprotein receptor (ASGPR) H2a (32). This glycoprotein is expressed naturally in hepatocytes as a membrane precursor, which undergoes efficient cleavage producing a 35-kDa secreted form. When H2a is expressed in other cell lines such as HEK 293, the membrane precursor is inefficiently cleaved, being retained in the ER, targeted to the ER-derived quality control compartment (ERQC), and then degraded by ERAD (32, 33). We further showed that ManIA is indeed involved in glycoprotein folding quality control, acting in parallel with ERManI and the EDEMs (23). Due to this surprising result, we examined possible roles of ManIB and ManIC in glycoprotein quality control in comparison with ManIA. In a pulse chase experiment, we confirmed that the overexpression of ManIA leads to a significantly increased degradation of H2a (Fig. 3A). Note the presence of a faster migrating underglycosylated species in the pulse. In the chase, the fully glycosylated species migrates faster due to mannose trimming, whereas the underglycosylated species is degraded. In contrast, the overexpression of ManIB and ManIC had almost no effect on the degradation of this substrate (Fig. 3A, B, E). Knockdown of ManIA strongly inhibited the degradation of H2a, and led to partial inhibition of mannose trimming, resulting in a slower migration in the chase compared to the control. Instead, there was almost no effect of the knockdown of ManIB or ManIC (Fig. 3C-F). The results suggest that these enzymes do not have redundant roles and that ManIB and ManIC are not involved in quality control of H2a.

**Figure 3.**
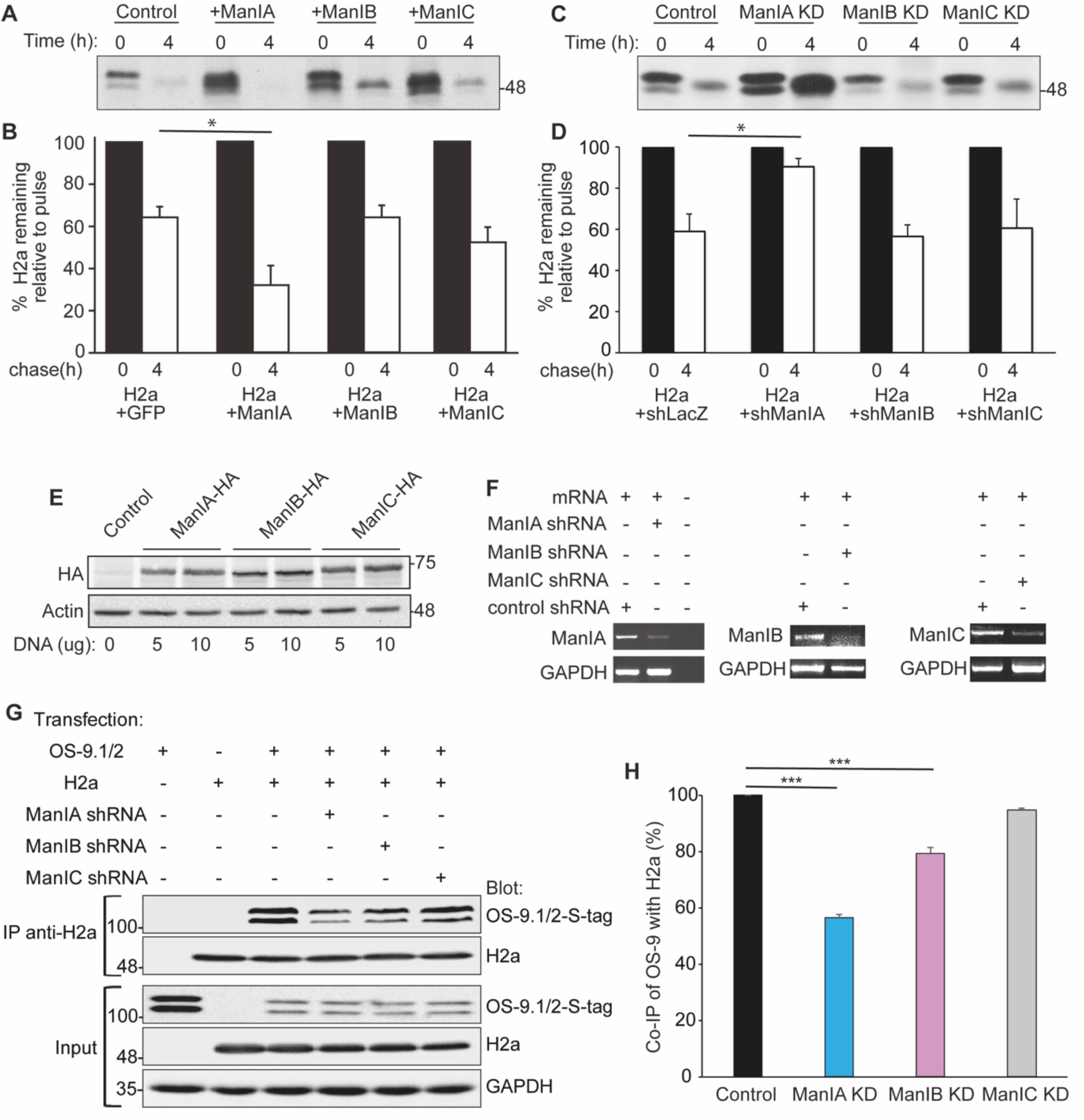
Overexpression of ManIA led to a significantly increased degradation of H2a, whereas its knockdown inhibited its degradation and its association with OS-9. ManIB and ManIC had almost no effect. **(A-D)** HEK 293 cells were co-transfected with H2a and ManIA-HA or ManIB-HA or ManIC-HA or control GFP (A), or with a pSUPER encoding anti-ManIA/B/C shRNA or with a pSUPER plasmid encoding control anti-lacZ shRNA (C). 48 hours post-transfection the cells were pulse-labeled for 20 min with [^35^S] Cys and chased for 0 h or 4 h in complete medium. Shown is a fluorography of labeled H2a, immunoprecipitated from cell lysates and run on SDS-PAGE. The graphs show the percentage of H2a remaining after chase relative to the pulse, from phosphor imager quantitation of the gels, average of three independent experiments ±SD (B, D). **(E)** An immunoblot of anti-HA antibody following overexpression of ManIA-HA, ManIB-HA or ManIC-HA, using 5 or 10 ug of DNA. **(F)** HEK 293 cells were transfected with a pSUPER plasmid encoding control anti-lacZ or anti-ManIA, anti-ManIB or anti-ManIC shRNA as indicated. RNA was extracted 48 h post-transfection and used for RT-PCR with primers for ManIA mRNA, ManIB mRNA, ManIC mRNA (top panels, as indicated) or GAPDH (bottom panels). No template was present in the control PCR reaction in lane 3. **(G)** HEK 293 cells were transfected with OS-9, H2a, and shRNA for ManIA, IB or IC. 48 hours post-transfection, the cells were lysed with buffer A with protease inhibitors. The lysates were incubated with anti-H2a antibody and protein A Sepharose beads. 10% of total lysates and the precipitates were run on a 10% SDS-PAGE and immunoblotted with anti-S-tag (OS-9) and anti-H2a antibodies. **(H)** The levels of OS-9 in the immunoprecipitate were normalized with OS-9 and H2a in the total lysates and compared with the control (=100). Average of three independent experiments ±SD.

According to the model of ERAD substrate targeting, ERAD-associated lectins OS-9 and XTP3-B, which have high affinity for glycoproteins after mannose trimming, trap the terminally misfolded glycoproteins inside the ERQC and facilitate the retrotranslocation (4, 10). Knockdown of the mannosidases may influence the mannose trimming and hence the association of ERAD substrates with the lectins. To examine the association between H2a and the lectin OS-9, we performed a co-immunoprecipitation experiment. HEK 293 cells transiently expressing H2a and S-tagged-OS-9.1/.2 together with or without shRNA for ManIA, ManIB or ManIC were lysed and immunoprecipitated with anti-H2a antibodies. The immunoprecipitate and 10% of total input lysates were immunoblotted to detect OS-9 and H2a. The relative amount of co-precipitated OS-9 decreased significantly upon knockdown of ManIA by approximately two-fold, while the effect of the knockdown of ManIB decreased the coimmunoprecipitation (coIP) by only 20% and ManIC had no effect (Fig. 3G, H).

### ManIA, ManIB and ManIC participate in the targeting to ERAD of NHK and BACE-I-476

Although ManIB and ManIC do not appear to participate in the quality control of H2a, we wondered whether they have substrate preference and might have a role in targeting other substrates to ERAD. We analyzed two other established ERAD substrates, NHK and a mutant (BACE-I-476) of the protein BACE, β-site APP-cleaving enzyme, which is retained in the ER (34). Pulse-chase analysis showed that the overexpression of each of the three enzymes separately or simultaneously accelerates the degradation of both NHK and BACE-I-476, compared to the control (Fig. 4A-D). The results show that at basal conditions, NHK was slowly degraded and after a 6-hour chase, there was a reduction by only roughly 10% in the amount of the substrate relative to the pulse. The overexpression of these enzymes had a significant effect, causing a reduction by about 40%-50% when being singly overexpressed and 60% following the overexpression of the three enzymes together. ManIA overexpression had the most significant effect on NHK compared to the other two enzymes overexpressed individually, whereas ManIB had the least effect. In contrast, ManIB had the strongest effect on BACE-I-476 (Fig. 4C, D). Upon knockdown of these enzymes, we observed an inhibition of the degradation of NHK. There was actually an increase in the amount of the substrate detected in the chase periods (Fig. 4E, F). An increase in labeling after short chases is common in pulse-chase analysis of relatively stable proteins, because of the incomplete chase of remaining radioactive precursor inside the cells. The results obtained with BACE-I-476 were similar to those of NHK, detecting an increase in the amount of the substrate following the knockdown of these enzymes individually or in combination after chase, compared to a small reduction in the amount of the substrate in the control (Fig. 4G, H). ManIC knockdown had the strongest effect on NHK, whereas ManIB knockdown had the most significant effect on BACE-I-476.

**Figure 4.**
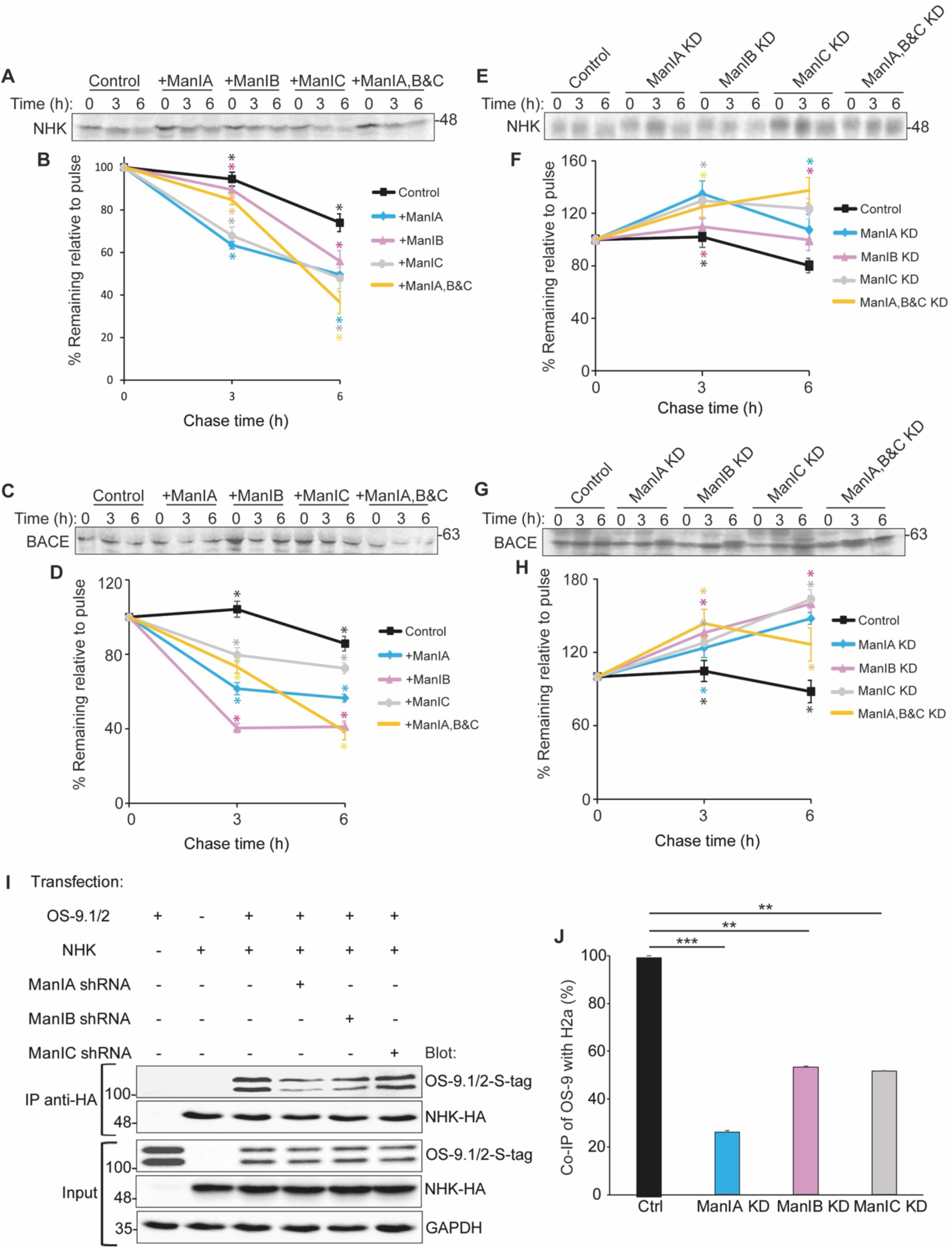
Overexpression of ManIA, ManIB and ManIC accelerates the degradation of the ERAD substrates NHK and BACE-I-476, while their knockdown has an inhibiting effect in degradation and association with OS-9. HEK 293 cells were co-transfected with either NHK **(A, E)** or BACE-I-476 **(C, G)** together with control GFP or ManIA-HA, ManIB-HA, ManIC-HA, separately (lanes 4-12) or simultaneously (lanes 13-15) **(A, C)**, or with a pSUPER plasmid encoding control anti-lacZ shRNA or a pSUPER encoding anti-ManIA/B/C shRNA separately (lanes 4-12) or simultaneously (lanes 13-15) **(E, G)**. 48 hours post-transfection, the cells were pulse-labeled for 20 min with [^35^S]Cys and chased for 0 h, 3 h or 6 h in complete medium. Shown is a fluorography of labeled NHK **(A, E)** and BACE-I-476 **(C, G)**, immunoprecipitated from cell lysates and run on SDS-PAGE. The graphs show the percentage of NHK **(B, F)** and BACE-I-476 **(D, H)** remaining after chase relative to the pulse, from phosphor imager quantitation of the gels, average of three independent experiments ±SD. P-values are of each chase vs. pulse. **(I)** HEK 293 cells were transfected with OS-9, NHK-HA, and shRNA for ManIA, IB, IC or control anti-lacZ. 48 hours post-transfection, the cells were lysed with buffer A with protease inhibitor. The lysates were incubated with anti-HA antibody and protein A Sepharose beads. 10% of total lysates (input) and the precipitates (IP) were run on a 10% SDS-PAGE and immunoblotted with anti-S-tag (OS-9) and anti-HA antibodies. **(J)** OS-9 in the immunoprecipitate was normalized with OS-9 and NHK in 10% of the total lysates and standardized with the control (=100). Average of three independent experiments ±SD.

Altogether, the results show that all 3 mannosidases appear to be involved in targeting to ERAD of at least two different ERAD substrate glycoproteins and indicate a substrate preference of each enzyme. It is also noteworthy that there is a lack of complementarity in the roles of the 3 enzymes, as both the overexpression or knockdown of all simultaneously failed to show a significant additive effect.

We also tested the effect of knockdown of the mannosidases on the coIP of OS-9. Although we could not detect measurable coIP with BACE-I-476, as we did in the case of NHK. Knockdown of any of the three mannosidases significantly decreased the coIP of OS-9 with NHK, with ManIA causing the strongest effect (Fig. 4I, J).

### Effects of knockout of the mannosidases on ERAD

To achieve higher depletion and evaluate the long-term effect of the absence of each of the mannosidases on the degradation of different ERAD substrates, we generated CRISPR/Cas9 knockout HEK 293 cell lines (Fig. 5A) and transfected them with either H2a (Fig. 5B, C), NHK (Fig. 5D, E) or BACE (Fig. 5F, G). ManIA knockout caused accumulation of all three substrates after 24h of their expression. In contrast, knockout of either ManIB or ManIC affected significantly NHK and BACE (∼40% increase) with only a minor effect on H2a. Although we cannot exclude effects on other processes, such as protein synthesis, the observed accumulation is most likely due to impaired degradation, as these results are consistent with the effects observed by knockdown and pulse chase analysis. Together, these findings strengthen the notion that all three mannosidases contribute to protein quality control and targeting of substrates to ERAD, while exhibiting distinct preferences for different ERAD substrates.

**Figure 5.**
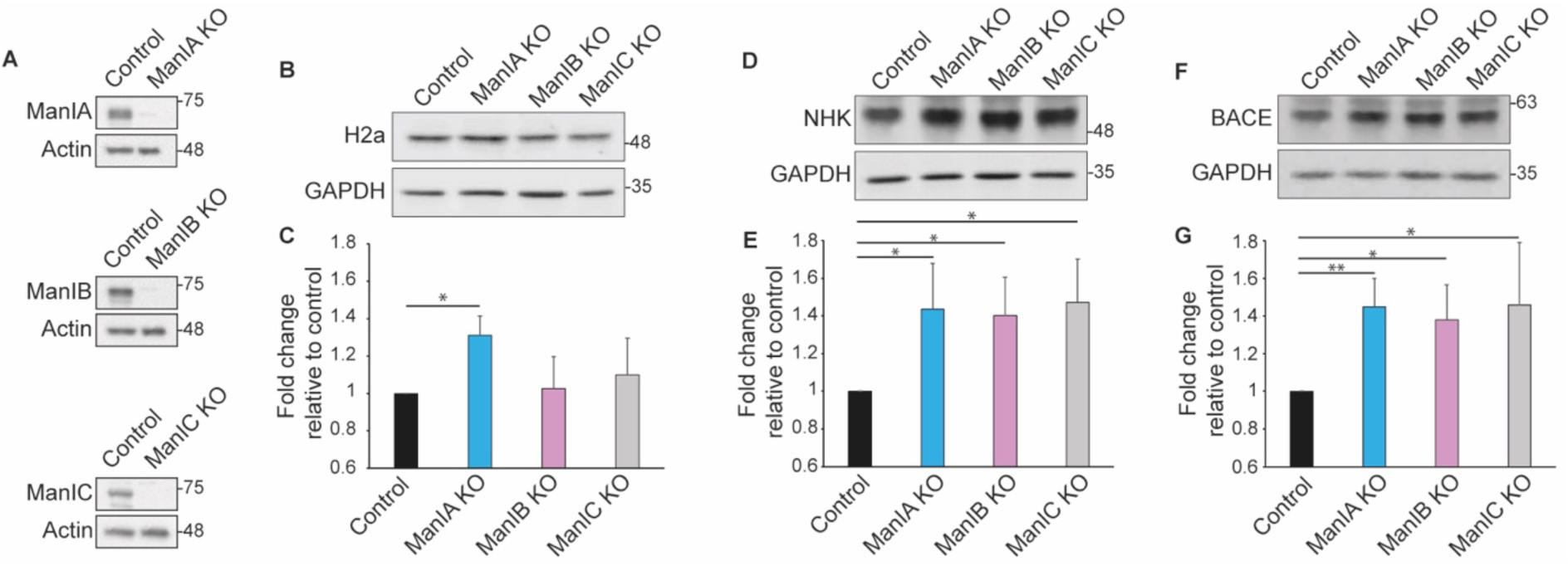
Knockouts of ManIB and ManIC have an inhibiting effect on the degradation of NHK and BACE, while ManIA knockout also affects the degradation of H2a. **(A)** Lysates of HEK 293 cells and ManIA, ManIB and ManIC HEK 293 knockout cells were run on 10% SDS-PAGE and immunoblotted with anti-ManIA, anti-ManIB or anti-ManIC antibodies. **(B-G)** HEK 293 cells or ManIA, ManIB or ManIC HEK 293 knockout cells were transfected with H2a **(B)** or NHK-HA **(D)** or BACE-HA **(F)**. 24h post-transfection the cells were lysed and the lysates were run on 10% SDS-PAGE and immunoblotted with anti-H2a or anti-HA antibodies. **(C, E, G)** The graphs show the fold change in each substrate relative to control. Average of three independent experiments ±SD.

### ManIA, ManIB, or ManIC modulate cell surface expression of a glycoprotein that matures through the Golgi and traffics to the plasma membrane

To investigate the role of these mannosidases in the quality control and maturation of glycoproteins that partially escape the ER, we utilized the asialoglycoprotein receptor H1 subunit as a model substrate. H1 is known to be able to exit the ER and reach the cell surface to a certain extent even when unassembled (32). HEK 293 cells were co-transfected with H1 and pSUPER encoding shRNA for ManIA, ManIB, or ManIC separately or simultaneously. At 48 h post-transfection, cells were incubated with the membrane-impermeable sulfo-succinimidyl-6-biotin to label surface proteins, followed by treatment with ammonium chloride to inactivate the reagent. The cells were then lysed with buffer A, and lysates were subjected to pull-down assays using streptavidin beads to isolate biotinylated proteins that reached the cell surface. Samples of the total lysates (Input) and the pull-down fractions were run on 10% SDS-PAGE and immunoblotted for H1 and GAPDH. As expected, quantification of the blots revealed that knockdown of ManIA, ManIB, ManIC or combined led to an increase in total H1 levels, indicating a likely inhibition of its degradation (Fig. 6A, B). On the other hand, the processing and maturation of H1, measured by the levels of a slower migrating endo H resistant band (Suppl. Fig. 1), was surprisingly not affected (Fig. 6C). As expected, only processed H1 reached the cell surface and appeared in the streptavidin pulldown. The cell surface expression of H1 was significantly increased by individual knockdowns of each enzyme, but not by their combination. Together, these results suggest that all 3 enzymes have an important role in the quality control of H1, but that they are redundant in its maturation and surface expression. However, at least one of the three must be present at sufficient levels for efficient H1 delivery to the cell surface.

**Figure 6.**
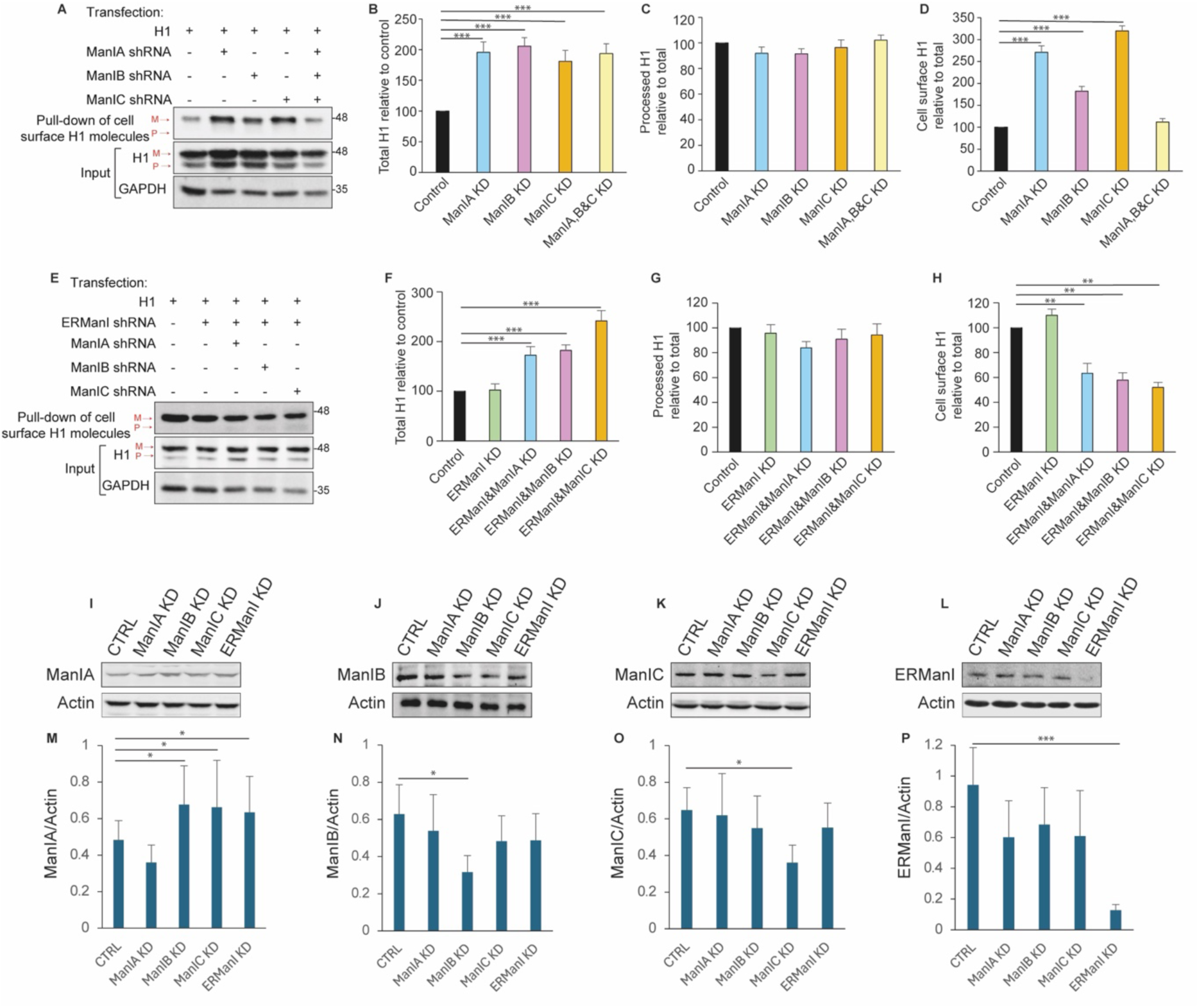
Combined knockdown of ManIA, ManIB, or ManIC inhibits the surface expression of asialoglycoprotein receptor H1. **(A, E)** HEK 293 cells were co-transfected with H1 and pSUPER encoding shRNA for ManIA, ManIB, or ManIC separately or simultaneously, or in combination with shRNA for ERManI (E). At 48 h post-transfection, the cells were incubated with sulfo succinimidyl-6-biotin to label surface proteins, followed by treatment with ammonium chloride to inactivate the reagent. Cells were then lysed and the lysates were subjected to pull-down assays using streptavidin beads to isolate biotinylated proteins that reached the cell surface. 10% of the total lysates (Input) and the pull-down samples were run on 10% SDS-PAGE and immunoblotted with anti-H1 and anti-GAPDH antibodies. The arrows indicate the migration of precursor H1 (P) and mature processed H1 (M). The blots were quantified using ImageJ, and the graphs for the experiment in (A): (B-D), or in (E): (F-H), represent the total H1 (B, F), processed H1 (C, G), and cell-surface H1 (D, H) relative to control, as the average of three independent experiments ± SD. **(I-L)** The lysates of cells from (A and E) were run on 10% SDS-PAGE and immunoblotted with anti-ManIA (I), ManIB (J), ManIC (K) or ERManI (L) antibodies. **(M-P)** The graphs represent the level of expression of ManIA (M), ManIB (N), ManIC (O) or ERManI (P), normalized by actin.

To better understand the participation of the different Class 1 α-mannosidases, ManIA, ManIB or ManIC were knocked down in combination with knockdown of ERManI. Combined knockdown of ERManI and ManIA, ManIB or especially ManIC accumulated synergistically the total amount of H1, suggesting a combined activity in their quality control targeting to ERAD (Fig. 6E, F). In contrast, H1 maturation was not affected (Fig. 6E, G) and H1 cell surface expression was significantly reduced (Fig. 6E, H). This suggests that there is redundancy in the roles of these enzymes in maturation, but that there is a synergy in the requirement of ERManI together with either ManIA, ManIB or ManIC for cell surface expression.

The levels of ManIB or ManIC were not affected by knockdown of the other enzymes (Fig. 6J, K, N, O). However, ManIA expression was considerably increased (Fig. 6I, M), suggesting a compensatory effect, which could explain the lack of effect on overall H1 maturation (Fig. 6C, G). Knockdown of any of the other enzymes caused some moderate concomitant reduction in ERManI levels (Fig. 6L, P), which might exacerbate the effect of the combined knockdown in interfering with H1 ERAD and with its cell surface expression.

We also assessed the effect of combined knockdown of ManIA, ManIB or ManIC with knockdown of the remaining Class 1 α-mannosidases, EDEM1, EDEM2 or EDEM3. The result was similar as with ERManI, with an increase in the total amount of H1 and little effect on maturation, except for a small and unexpected increase in processing when knockdown of ManIA, ManIB or ManIC was combined with the knockdown of EDEM1 (Suppl. Fig. S2). The cell surface expression of H1 was not significantly affected, despite the increase in total H1. This suggests that similarly to ERManI, the EDEMs have a combined activity with ManIA, ManIB and ManIC in targeting to ERAD, that they are redundant in maturation and that they act synergistically in facilitating cell surface expression.

The levels of the EDEMs were increased in some cases as a compensatory effect for the knockdown of the other enzymes. This was especially noticeable for EDEM3, the expression of which was substantially increased upon knockdown of all other enzymes except for EDEM1 (Suppl. Fig S2 Q, R).

### ManIA, ManIB and ManIC have a preference for processing of the N-linked sugar chains on a native compared to a denatured substrate glycoprotein

Our results with ERAD substrates and with H1 indicate that all 3 mannosidases ManIA, ManIB and ManIC have roles in both glycoprotein quality control and maturation. Given the significant effect of the knockdown of the individual mannosidases on the quality control of H1, but a lack of effect or redundancy in H1 maturation (Fig. 6), we tried to better understand their roles by measuring their activity in vitro on a native or denatured high mannose glycoprotein, as models of a properly folded glycoprotein or a misfolded substrate. High-performance liquid chromatography (HPLC) analysis enables the evaluation of the effects of the mannosidases on the structure of N-linked sugar chains. We treated M. rosenbergii vitellogenin, which we had used previously as a model glycoprotein because it bears a series of high-mannose N-linked oligosaccharides(8), with each of the mannosidases. We then quantified after HPLC the presence of different species in the high mannose oligosaccharide mix released from the glycoprotein with endo H. The fully mannosylated Man_9_GlcNAc (M9) is illustrated in Fig. 7A. When the glycoprotein is at its native state, all three mannosidases showed significant mannose trimming activity, although to different extents for each mannosidase (Fig. 7 B-E). ManIA showed the highest activity in trimming down from M9 and M8 to the M5 species (lacking all α1,2-linked mannose residues (Fig. 7 C, J, K). ManIB trimmed down preferentially to M6 rather than to M5. ManIC showed much less activity in using M9 as a substrate but trimmed more efficiently from M8 and M7 to M6 and M5.

**Figure 7.**
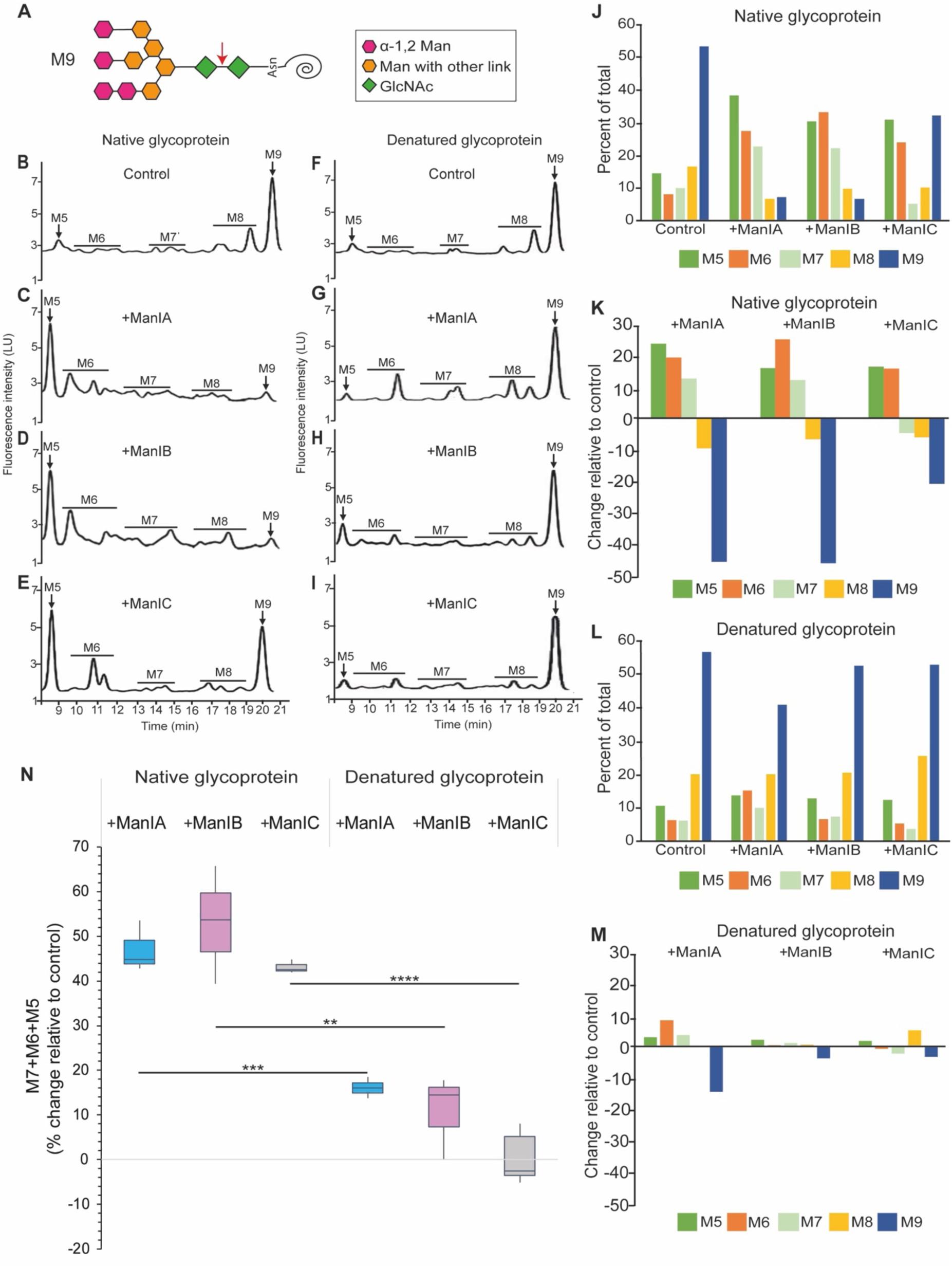
High-performance liquid chromatography (HPLC) analysis of the effects of ManIA, ManIB or ManIC on the N-linked glycans of a native glycoprotein and its denatured form. **(A)** Scheme of a high mannose N-linked glycan, possessing all 9 mannose residues (M9). The arrow indicates the location of the Endoglycosidase H (Endo H) cleavage site. (**B-I)** NP profiles of (B, F) Control, (C, G) ManIA-HA, (D, H) ManIB-HA and (E, I) ManIC-HA 2AB labeled Endo H treated glycans on (B-E) native and (F-I) denatured glycoprotein. HPLC chromatograms of an oligosaccharide mix, released from M. rosenbergii vitellogenin, after treatment in vitro with ManIA-HA, ManIB-HA or ManIC-HA expressed in HEK 293 cells and immunoprecipitated with anti-HA. For the control, immunoprecipitated ManIB was inactivated by boiling prior to the incubation with the glycoprotein. A representative of three independent experiments is presented. **(J, L)** The chromatograms were quantified, and the amount of each species was plotted as percent of total oligosaccharides for native (J) or denatured glycoprotein (L). **(K, M)** The relative change in each species compared to the control was calculated and plotted. **(N)** The total relative change in the sum of M5, M6, and M7 species in the presence of each mannosidase using native or denatured vitellogenin. Data from three independent experiments are presented as box-and-whisker plots. The center line indicates the median, the box extends from the first to the third quartile, and the whiskers indicate the minimum and maximum values.

In contrast to the native glycoprotein, the processing of the sugar chains from denatured vitellogenin was much less efficient for all 3 enzymes, with the highest activity for ManIA and the lowest for ManIC. Noteworthy, ManIC processing was mostly from M9 to M8, with negligible further trimming (Fig. 7I, L, M). The extent of extensive trimming can be better illustrated by the change in the sum of the M5, M6 and M7 species relative to the control (Fig. 7N). All three mannosidases have significantly higher activity on the native glycoprotein compared to the denatured form. ManIB showed the highest activity in trimming down to M5-M7 on the native glycoprotein, compared to ManIA and ManIC (Fig. 7N). Although all three enzymes showed very low activity on the denatured glycoprotein, ManIC was the lowest.

Together, the results indicate that there are small differences in the activity and extent of processing of a substrate in a particular conformation by ManIA, ManIB or ManIC, but there is a major increase in the activity of all three enzymes on a native glycoprotein compared to its denatured form. This result is surprising, as it is the opposite of what was observed with ERManI, EDEM1 and EDEM2, which have a much higher activity on a denatured substrate (8).

## Discussion

Despite the long-recognized importance of α-1,2 mannose trimming in determining the fate of N-linked glycoproteins, both in quality control targeting to degradation (ERAD) and in their Golgi maturation, it has remained unclear why mammalian cells require seven different Class I α-1,2 mannosidases and whether these enzymes perform distinct functions (35, 36). It was originally assumed that ERManI and the EDEMs function primarily in quality control, whereas the three “Golgi” mannosidases, ManIA, ManIB, and ManIC, are dedicated to glycoprotein maturation (19, 36). Our findings challenge this model by demonstrating that ManIA and ManIB are largely absent from the Golgi, while ManIC is only partially localized there. Moreover, all three enzymes contribute to both glycoprotein quality control and maturation.

Immunofluorescence and live-cell imaging revealed that these mannosidases predominantly localize to punctate vesicular structures, colocalizing with Rab1a and with only partial colocalization of ManIC and not of ManIB with conventional Golgi and ERGIC markers. The partial overlap of ManIB- and ManIC-containing vesicles with ManIA-positive structures (Fig. 1), together with their BFA-insensitivity and their ER-like density determined by equilibrium sedimentation gradients (Fig. 2), suggest that, similarly to ManIA, ManIB and in part ManIC also reside in QCVs. Moreover, both ManIB- and ManIC-containing vesicles showed high mobility, which was reduced by expression of dominant-negative Sar1, indicating that their dynamics depends, at least in part, on Sar1-dependent trafficking, as had been observed for QCVs (4). We previously characterized QCVs as COPII-dependent vesicles with an ER-like density, although COPII coat components themselves were not detected on these structures. ERManI and ManIA were found to localize mainly to QCVs, where they encounter their glycoprotein substrates, possibly through transient fusion events with vesicular or organellar compartments carrying the cargo (4, 23). EDEM1 also resides mostly in vesicles (37), possibly in QCVs. Velocity sedimentation gradients further showed that, despite their high sequence homology, ManIA, ManIB and ManIC segregate into vesicular populations that are surprisingly distinct (Fig. 2). ManIB localized to vesicles of intermediate density and size (∼120–150 nm), whereas ManIC was associated with significantly lighter and smaller vesicles (∼50– 80 nm). In contrast, ManIA was present in vesicular structures of intermediate density (23) but substantially larger size (>200 nm) (Fig. 2). These distinct localizations may reflect interactions with specific Golgi or vesicular proteins that determine enzyme trafficking and function (38). Such spatial partitioning possibly underlies the substrate-specific ERAD roles observed in this study (9). ManIA broadly influenced the degradation of H2a, NHK, and BACE, whereas ManIB and ManIC selectively affected NHK and BACE degradation without significantly altering H2a turnover (Fig. 3–5). This specificity was further supported by co-immunoprecipitation experiments showing that ManIA knockdown reduced the interaction between H2a and the ERAD lectin OS-9, while knockdown of ManIB or ManIC had no detectable effect (Fig. 3). Unlike NHK, H2a undergoes true ER retention rather than ER exit and retrieval (32). The trimming of α-1,2 mannose residues on the N-glycans of H2a was shown to be essential for its delivery to ERAD, with a particularly strong dependence on ERManI (5, 6).

Our analysis of the asialoglycoprotein receptor H1 subunit provides further insight into how these enzymes regulate the fate of glycoproteins that partially escape the ER, mature through the Golgi, and reach the cell surface. Knockdown of ManIA, ManIB, or ManIC individually increased total H1 levels, indicating that all three enzymes contribute to H1 quality control and targeting for degradation. Surprisingly, however, neither individual nor combined knockdown of all three mannosidases, produced a detectable defect in H1 maturation, even in combination with knockdown of ERManI or the EDEMs. These findings suggest an unexpectedly robust redundancy among Class I α-1,2 mannosidases in glycan processing. The compensatory increase in ManIA and EDEM3 expression following depletion of other mannosidases (Fig. 6 and Fig. S2) complicates straightforward interpretation of knockdown phenotypes, but points to a particularly important buffering role for these enzymes. Consistent with this interpretation, previous studies demonstrated that complex and hybrid glycan structures are abolished only after simultaneous knockout of four mannosidases - ERManI, ManIA, ManIB, and ManIC (39, 40), further supporting extensive functional redundancy in glycan processing.

At the same time, the marked inhibition of cell-surface expression of H1 molecules that accumulate and mature following simultaneous depletion of ManIA, ManIB, and ManIC suggests that the activity of at least one of these enzymes is required for a late quality-control checkpoint (Fig. 6). This requirement may reflect the need for proper glycan processing of one or more factors essential for Golgi-to-cell surface trafficking.

Our HPLC analysis of N-linked glycans revealed an unexpected preference of ManIA, ManIB, and ManIC for native rather than denatured (misfolded) glycoprotein molecules, at least for the vitellogenin model glycoprotein (Fig. 7). In contrast, ERManI, EDEM1, and EDEM2 preferentially act on misfolded substrates (8), suggesting that the glycoprotein “quality-control timer” is considerably more sophisticated than originally thought (3, 41, 42) (Fig. 8). In this model, ERManI and the EDEMs act relatively slowly on partially folded intermediates, thereby extending the window for productive folding, while rapidly trimming terminally misfolded proteins, likely near OS-9-containing ERAD complexes, to commit them to degradation. Conversely, ManIA, ManIB, and ManIC may act coordinately but more slowly on misfolded glycoproteins, allowing their retrograde transfer to the ER and subsequently to the ERQC through interactions with ER-Golgi lectins such as VIPL, VIP36, and ERGIC53 (43). In contrast, these enzymes would rapidly trim properly folded glycoproteins, preventing lectin-mediated retention or retrieval and thereby promoting their release for forward trafficking and maturation in the Golgi (16).

**Figure 8.**
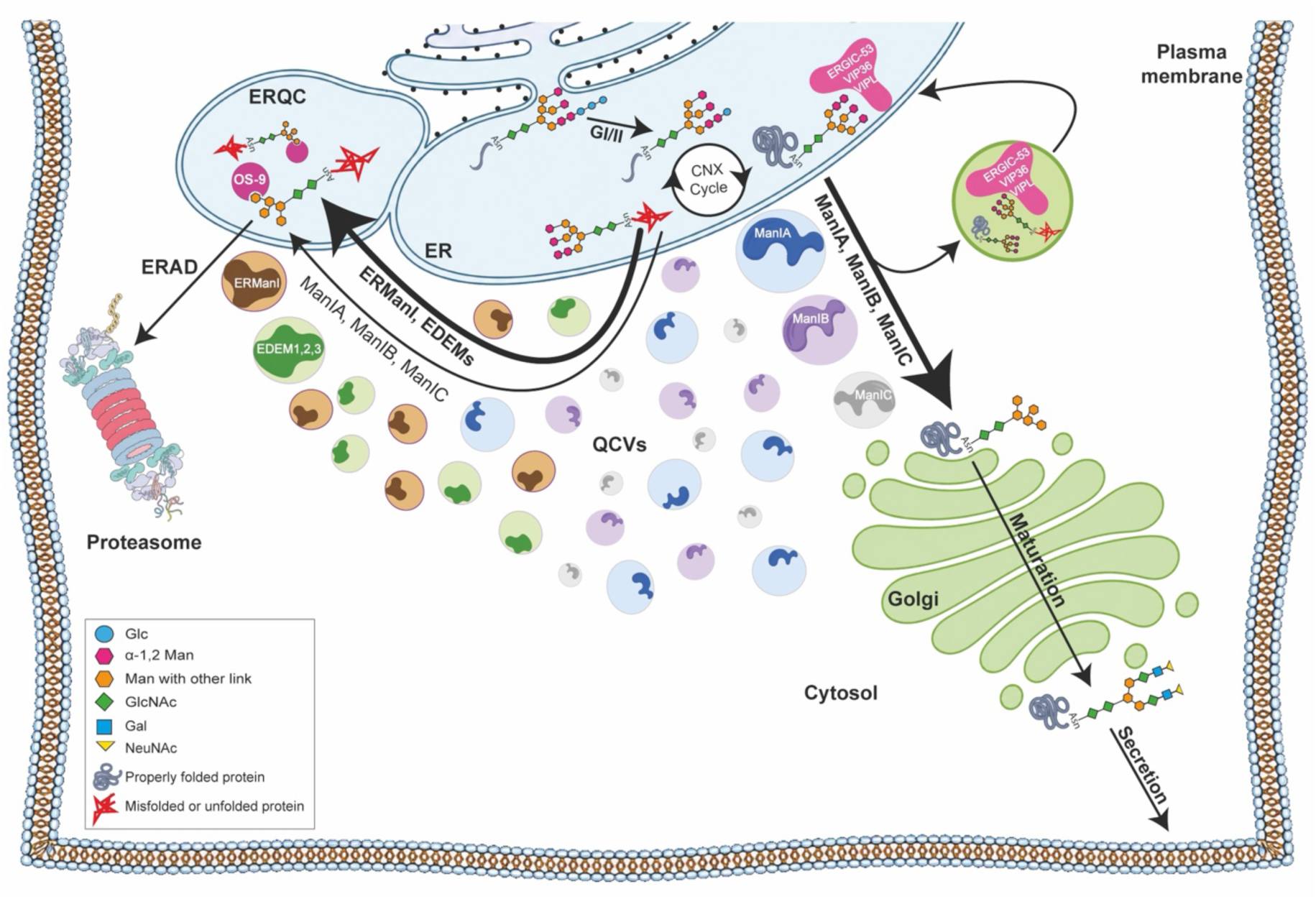
A model for glycoprotein quality control. Newly synthesized glycoproteins enter the calnexin (CNX) folding cycle in the ER, where they can either achieve a properly folded conformation or become terminally misfolded. The Class I α-1,2 mannosidases are localized within distinct quality control vesicles (QCVs), and likely process glycoprotein substrates through transient interactions or fusion events between QCVs and vesicular or organellar compartments carrying these cargoes. In this model, ManIA, ManIB, and ManIC preferentially act on properly folded glycoproteins, rapidly trimming their mannose residues, although they can also process misfolded substrates at a slower rate. Folding intermediates bearing untrimmed high-mannose glycans can be recognized by the lectins ERGIC-53, VIPL, and VIP36, which may mediate retrieval to the ER. Mannose trimming by ManIA, ManIB, and ManIC prevents binding to these lectins, facilitating release of properly folded glycoproteins for forward trafficking, Golgi maturation and subsequent transport to the cell surface or other final destinations. In contrast, ERManI and the EDEMs preferentially process misfolded glycoproteins, accelerating mannose trimming to generate glycan structures recognized by the ERAD lectin OS-9 in the ER quality control compartment (ERQC). Recognition by OS-9 commits terminally misfolded glycoproteins to ER-associated degradation (ERAD).

## Acknowledgments

We would like to thank Navit Ogen-Shtern and Ron Benyair for initial work related to this study, Dan Peer for advice and Nicole Gorokhovsky for help with DLS, Vered Holdengreber for help with EM and Eran Bacharach for plasmids. GPT-5.5 was used to polish parts of the text. All AI-assisted text was reviewed and revised by the authors to ensure accuracy and clarity of meaning. Work was supported by grant 2577/20 from the Israel Science Foundation (GZL).

## Data Availability

The data that support the findings of this study are available in the Materials and Methods, Results, and/or Supplemental Material of this article.

## Author Contributions

Conceptualization: GZL. Investigation: HS, MS, EA. Methodology: IK. Supervision: GZL. Writing: HS, GZL. Review and editing: HS, MS, EA, IK, GZL.

## References

1. Aebi, M., Bernasconi, R., Clerc, S., and Molinari, M. (2010) N-glycan structures: recognition and processing in the ER. Trends in biochemical sciences 35, 74–82

2. Caramelo, J. J., and Parodi, A. J. (2015) A sweet code for glycoprotein folding. FEBS letters 589, 3379–3387

3. Lederkremer, G. Z. (2009) Glycoprotein folding, quality control and ER-associated degradation. Curr Opin Struct Biol 19, 515–523

4. Benyair, R., Ogen-Shtern, N., Mazkereth, N., Shai, B., Ehrlich, M., and Lederkremer, G. Z. (2015) Mammalian ER mannosidase I resides in quality control vesicles, where it encounters its glycoprotein substrates. Mol Biol Cell 26, 172–184

5. Frenkel, Z., Gregory, W., Kornfeld, S., and Lederkremer, G. Z. (2003) Endoplasmic reticulum-associated degradation of mammalian glycoproteins involves sugar chain trimming to Man6-5GlcNAc2. The Journal of biological chemistry 278, 34119–34124

6. Avezov, E., Frenkel, Z., Ehrlich, M., Herscovics, A., and Lederkremer, G. Z. (2008) Endoplasmic reticulum (ER) mannosidase I is compartmentalized and required for N-glycan trimming to Man5-6GlcNAc2 in glycoprotein ER-associated degradation. Mol Biol Cell 19, 216–225

7. Olivari, S., and Molinari, M. (2007) Glycoprotein folding and the role of EDEM1, EDEM2 and EDEM3 in degradation of folding-defective glycoproteins. FEBS Lett 581, 3658–3664

8. Shenkman, M., Ron, E., Yehuda, R., Benyair, R., Khalaila, I., and Lederkremer, G. Z. (2018) Mannosidase activity of EDEM1 and EDEM2 depends on an unfolded state of their glycoprotein substrates. Communications biology 1, 172

9. Shenkman, M., and Lederkremer, G. Z. (2019) Compartmentalization and Selective Tagging for Disposal of Misfolded Glycoproteins. Trends Biochem Sci 44, 827–836

10. Mikami, K., Yamaguchi, D., Tateno, H., Hu, D., Qin, S. Y., Kawasaki, N., Yamada, M., Matsumoto, N., Hirabayashi, J., Ito, Y., and Yamamoto, K. (2010) The sugar-binding ability of human OS-9 and its involvement in ER-associated degradation. Glycobiology 20, 310–321

11. Groisman, B., Shenkman, M., Ron, E., and Lederkremer, G. Z. (2011) Mannose trimming is required for delivery of a glycoprotein from EDEM1 to XTP3-B and to late endoplasmic reticulum-associated degradation steps. The Journal of biological chemistry 286, 1292–1300

12. Ritter, C., and Helenius, A. (2000) Recognition of local glycoprotein misfolding by the ER folding sensor UDP-glucose:glycoprotein glucosyltransferase. Nature structural biology 7, 278–280

13. Caramelo, J. J., and Parodi, A. J. (2008) Getting in and out from calnexin/calreticulin cycles. The Journal of biological chemistry 283, 10221–10225

14. Lamriben, L., Graham, J. B., Adams, B. M., and Hebert, D. N. (2016) N-Glycan-based ER Molecular Chaperone and Protein Quality Control System: The Calnexin Binding Cycle. Traffic (Copenhagen, Denmark) 17, 308–326

15. Hosokawa, N., Kamiya, Y., Kamiya, D., Kato, K., and Nagata, K. (2009) Human OS-9, a lectin required for glycoprotein endoplasmic reticulum-associated degradation, recognizes mannose-trimmed N-glycans. J Biol Chem 284, 17061–17068

16. Kamiya, Y., Kamiya, D., Yamamoto, K., Nyfeler, B., Hauri, H. P., and Kato, K. (2008) Molecular basis of sugar recognition by the human L-type lectins ERGIC-53, VIPL, and VIP36. The Journal of biological chemistry 283, 1857–1861

17. Neve, E. P., Svensson, K., Fuxe, J., and Pettersson, R. F. (2003) VIPL, a VIP36-like membrane protein with a putative function in the export of glycoproteins from the endoplasmic reticulum. Experimental cell research 288, 70–83

18. Xiang, Y., Karaveg, K., and Moremen, K. W. (2016) Substrate recognition and catalysis by GH47 α-mannosidases involved in Asn-linked glycan maturation in the mammalian secretory pathway. Proceedings of the National Academy of Sciences of the United States of America 113, E7890–e7899

19. Herscovics, A. (2001) Structure and function of Class I alpha 1,2-mannosidases involved in glycoprotein synthesis and endoplasmic reticulum quality control. Biochimie 83, 757–762

20. Lal, A., Pang, P., Kalelkar, S., Romero, P. A., Herscovics, A., and Moremen, K. W. (1998) Substrate specificities of recombinant murine Golgi alpha1, 2-mannosidases IA and IB and comparison with endoplasmic reticulum and Golgi processing alpha1,2-mannosidases. Glycobiology 8, 981–995

21. Hosokawa, N., You, Z., Tremblay, L. O., Nagata, K., and Herscovics, A. (2007) Stimulation of ERAD of misfolded null Hong Kong alpha1-antitrypsin by Golgi alpha1,2-mannosidases. Biochem Biophys Res Commun 362, 626–632

22. Demaretz, S., Seaayfan, E., Bakhos-Douaihy, D., Frachon, N., Kömhoff, M., and Laghmani, K. (2021) Golgi Alpha1,2-Mannosidase IA Promotes Efficient Endoplasmic Reticulum-Associated Degradation of NKCC2. Cells 11

23. Ogen-Shtern, N., Avezov, E., Shenkman, M., Benyair, R., and Lederkremer, G. Z. (2016) Mannosidase IA is in Quality Control Vesicles and Participates in Glycoprotein Targeting to ERAD. Journal of molecular biology 428, 3194–3205

24. Saad, H., Benyair, R., Mazor, T., and Lederkremer, G. Z. (2025) Helical charge distribution at the transmembrane-luminal interface determines subcellular localization. bioRxiv, 2025.2011.2012.688035

25. Shenkman, M., Ogen-Shtern, N., Patel, C., Saad, H., Groisman, B., Pasmanik-Chor, M., Schermann, S. M., Körner, R., and Lederkremer, G. Z. (2025) Oligosaccharyltransferase Is Involved in Targeting to ER-Associated Degradation. Cells 14

26. Sharma, N., Patel, C., Shenkman, M., Kessel, A., Ben-Tal, N., and Lederkremer, G. Z. (2021) The Sigma-1 receptor is an ER-localized type II membrane protein. The Journal of biological chemistry 297, 101299

27. Ogen-Shtern, N., Chang, C., Saad, H., Mazkereth, N., Patel, C., Shenkman, M., and Lederkremer, G. Z. (2023) COP I and II dependent trafficking controls ER-associated degradation in mammalian cells. iScience 26, 106232

28. Doms, R. W., Russ, G., and Yewdell, J. W. (1989) Brefeldin A redistributes resident and itinerant Golgi proteins to the endoplasmic reticulum. The Journal of cell biology 109, 61–72

29. Benyair, R., and Lederkremer, G. Z. (2016) Common fixation-permeabilization methods cause artifactual localization of a type II transmembrane protein. Microscopy (Oxf) 65, 517–521

30. Nuoffer, C., Davidson, H. W., Matteson, J., Meinkoth, J., and Balch, W. E. (1994) A GDP-bound of rab1 inhibits protein export from the endoplasmic reticulum and transport between Golgi compartments. The Journal of cell biology 125, 225–237

31. Ward, T. H., Polishchuk, R. S., Caplan, S., Hirschberg, K., and Lippincott-Schwartz, J. (2001) Maintenance of Golgi structure and function depends on the integrity of ER export. The Journal of cell biology 155, 557–570

32. Tolchinsky, S., Yuk, M. H., Ayalon, M., Lodish, H. F., and Lederkremer, G. Z. (1996) Membrane-bound versus secreted forms of human asialoglycoprotein receptor subunits. Role of a juxtamembrane pentapeptide. The Journal of biological chemistry 271, 14496–14503

33. Kamhi-Nesher, S., Shenkman, M., Tolchinsky, S., Fromm, S. V., Ehrlich, R., and Lederkremer, G. Z. (2001) A novel quality control compartment derived from the endoplasmic reticulum. Molecular biology of the cell 12, 1711–1723

34. Tanahashi, H., and Tabira, T. (2001) Three novel alternatively spliced isoforms of the human beta-site amyloid precursor protein cleaving enzyme (BACE) and their effect on amyloid beta-peptide production. Neuroscience letters 307, 9–12

35. Mast, S. W., and Moremen, K. W. (2006) Family 47 alpha-mannosidases in N-glycan processing. Methods in enzymology 415, 31–46

36. Moremen, K. W., and Molinari, M. (2006) N-linked glycan recognition and processing: the molecular basis of endoplasmic reticulum quality control. Current opinion in structural biology 16, 592–599

37. Calì, T., Galli, C., Olivari, S., and Molinari, M. (2008) Segregation and rapid turnover of EDEM1 by an autophagy-like mechanism modulates standard ERAD and folding activities. Biochem Biophys Res Commun 371, 405–410

38. Bhat, G., Hothpet, V. R., Lin, M. F., and Cheng, P. W. (2017) Shifted Golgi targeting of glycosyltransferases and α-mannosidase IA from giantin to GM130-GRASP65 results in formation of high mannose N-glycans in aggressive prostate cancer cells. Biochim Biophys Acta Gen Subj 1861, 2891–2901

39. Jin, Z. C., Kitajima, T., Dong, W., Huang, Y. F., Ren, W. W., Guan, F., Chiba, Y., Gao, X. D., and Fujita, M. (2018) Genetic disruption of multiple α1,2-mannosidases generates mammalian cells producing recombinant proteins with high-mannose-type N-glycans. J Biol Chem 293, 5572–5584

40. Ren, W. W., Jin, Z. C., Dong, W., Kitajima, T., Gao, X. D., and Fujita, M. (2019) Glycoengineering of HEK293 cells to produce high-mannose-type N-glycan structures. J Biochem 166, 245–258

41. Helenius, A. (1994) How N-Linked Oligosaccharides Affect Glycoprotein Folding in the Endoplasmic Reticulum. Mol Biol Cell 5, 253–265

42. Tannous, A., Pisoni, G. B., Hebert, D. N., and Molinari, M. (2014) N-linked sugar-regulated protein folding and quality control in the ER. Seminars in cell & developmental biology

43. Reiterer, V., Nyfeler, B., and Hauri, H. P. (2010) Role of the lectin VIP36 in post-ER quality control of human alpha1-antitrypsin. Traffic (Copenhagen, Denmark) 11, 1044–1055

