## Supplementary material for "Mannosidases IA, IB and IC are in segregated vesicular structures and involved in both glycoprotein quality control and maturation": Suppl. Fig.

**Supplementary figures:**

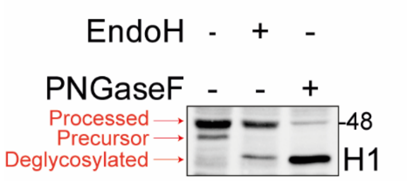


**Figure S1.** HEK 293 cells were transfected with H1. 24h post-transfection, the cells were lysed, and the lysates were subjected to enzymatic digestion with Endoglycosidase H (EndoH) or peptide N-glycosidase F (PNGaseF). H1 displays two distinct bands – a slower migrating processed form and a faster migrating precursor form.

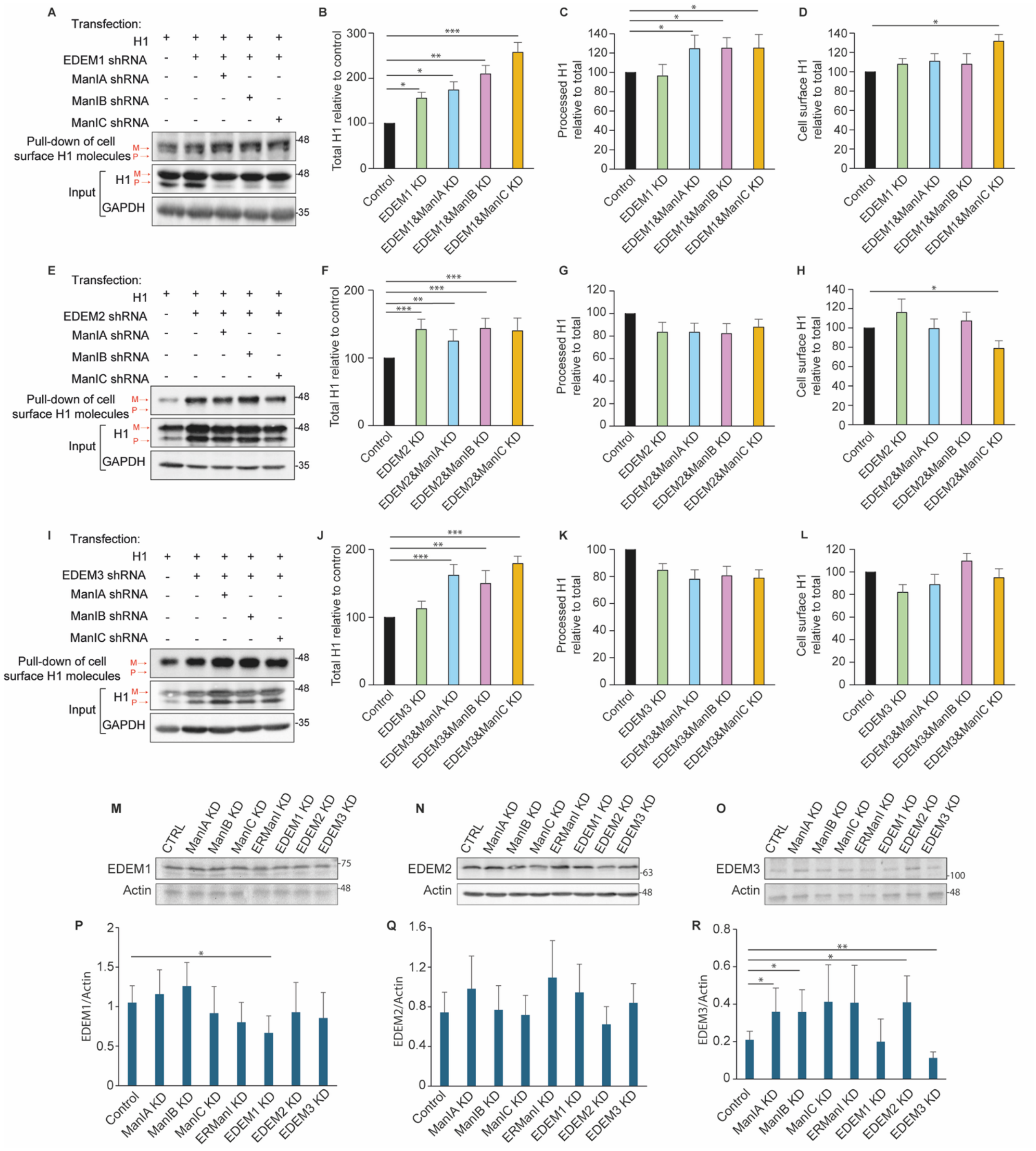


**Figure S2. Combined knockdown of ManIA, ManIB, or ManIC and EDEMs affects quality control of asialoglycoprotein receptor H1. (A, E, I)** HEK 293 cells were co-transfected with H1 and pSUPER encoding shRNA for ManIA, ManIB, or ManIC in combination with shRNA for EDEM1, EDEM2, or EDEM3. They were then processed as in Fig. 6 to isolate biotinylated proteins that reached the cell surface. The arrows indicate the migration of precursor H1 (P) and mature processed H1 (M). The blots were quantified using ImageJ, and the graphs for the experiment in (A): (B-D), in (E): (F-H), or in (I): (J-L) represent the total H1 (B, F, J), processed

H1 (C, G, K), and cell-surface H1 (D, H, L) relative to control as the average of three independent experiments ± SD. **(M-O)** The lysates of cells from (A, E, I) were run on 10% SDS-PAGE and immunoblotted with anti-EDEM1 (M), EDEM2 (N), or EDEM3 (O) antibodies. **(P-R)** The graphs represent the level of expression of EDEM1 (P), EDEM2 (Q), or EDEM3 (R), normalized by actin.

### Supplementary video legends:

**Video S1. Live-cell imaging of ManIB-mCherry and GalT-YFP.** Time-lapse live-cell imaging of NIH 3T3 cells co-expressing ManIB-mCherry and the Golgi marker GalT-YFP. Only the mCherry channel is shown. The movie spans 4 min 13 s. The time points T = 0, 98, and 192 s correspond to those presented in Fig. 2D right panel.

**Video S2. Live-cell imaging of ManIB-mCherry in cells expressing Sar1[T39N]-GFP.** Time-lapse live-cell imaging of NIH 3T3 cells co-expressing ManIB-mCherry and the dominant-negative Sar1[T39N]-GFP mutant. Only the mCherry channel is shown. The movie spans 5 min 03 s. The time points T = 0, 98, and 192 s correspond to those presented in Fig. 2E right panel.

**Video S3. Live-cell imaging of ManIC-mCherry and GalT-YFP.** Time-lapse live-cell imaging of NIH 3T3 cells co-expressing ManIC-mCherry and the Golgi marker GalT-YFP. Only the mCherry channel is shown. The movie spans 3 min 27 s. The time points T = 0, 98, and 192 s correspond to those presented in Fig. 2F right panel.

**Video S4. Live-cell imaging of ManIC-mCherry in cells expressing Sar1[T39N]-GFP.** Time-lapse live-cell imaging of NIH 3T3 cells co-expressing ManIC-mCherry and the dominant-negative Sar1[T39N]-GFP mutant. Only the mCherry channel is shown. The movie spans 3 min 17 s. The time points T = 0, 98, and 192 s correspond to those presented in Fig. 2G right panel.
